# A re-evaluation of Muller’s sheltering hypothesis for the evolution of sex chromosome gene content

**DOI:** 10.1101/2025.04.24.650552

**Authors:** Andrea Mrnjavac, Beatriz Vicoso, Tim Connallon

## Abstract

The first influential hypothesis for sex chromosome evolution was proposed in 1914 by H. J. Muller, who argued that once recombination was suppressed between the X and Y chromosomes, Y-linked genes become “sheltered” from selection, leading to accumulation of recessive loss-of-function (LOF) mutations and decay of Y-linked genes. The hypothesis fell out of favour in the 1970s because early mathematical models failed to support it and data on the dominance of lethal mutations were viewed as incompatible with the hypothesis. We reevaluate the main arguments against Muller’s hypothesis and find that they do not conclusively exclude a role for sheltering in sex chromosome evolution. By relaxing restrictive assumptions of earlier models, we show that sheltering promotes fixation of LOF mutations with sexually dimorphic fitness effects, resulting in decay of X-linked genes that are exclusively expressed by males and Y-linked genes that are primarily, though not necessarily exclusively, expressed by females. We further show that drift and other processes contributing to Y degeneration (i.e., selective interference and regulatory evolution) expand conditions of Y-linked gene loss by sheltering. The actual contribution of sheltering to sex chromosome evolution hinges upon the distribution of dominance and sex-specific fitness effects of LOF mutations, which we discuss.

## Introduction

Each new X and Y chromosome pair is undifferentiated, yet this initial state is typically transient (Charlesworth 1996; Bachtrog 2013; Bachtrog et al. 2014; for exceptions, see: Stöck et al. 2011; Kamiya et al. 2012; Vicoso et al. 2013). The suppression of recombination between the X and Y initiates a cascade of evolutionary changes that ultimately leads to differentiation and gene losses from the Y (Bachtrog 2013; Abbott et al. 2017; Furman et al. 2020), though recent studies on *Drosophila* and mammals also report gene losses from the X (Nozawa et al. 2016, 2021; Hughes et al. 2022). The X and Y also evolve differences in the *types* of genes that they carry, with the Y typically becoming enriched for genes involved in male fertility (Skaletsky et al. 2003; Bellot et al. 2014; Mahajan and Bachtrog 2017; Nozawa et al. 2021; Shaw and White 2022; Wei et al. 2024), and the X becoming enriched for genes preferentially expressed by females (‘female-biased genes’) and strongly deficient in genes primarily expressed by males (Kaiser and Ellegren 2006; Ellegren 2011; Meisel et al. 2012; Albritton et al. 2014; Papa et al. 2017; Foster et al. 2020; Hu et al. 2022; Lasne et al. 2023; Mora et al. 2024).

Several processes are thought to explain these patterns of sex chromosome gene content evolution. Y-linked gene losses are widely attributed to selective interference (i.e., Hill-Robertson effects), in which natural selection becomes overwhelmed by genetic drift in genomic regions that lack recombination (Felsenstein 1974; Charlesworth 1978; Bachtrog 2013; Charlesworth and Campos 2014). Under this view, the lack of crossing over between the X and Y hinders the evolutionary removal of mildly deleterious mutations and the fixation of beneficial mutations, resulting in a gradual decay of functional Y-linked genes, and retention of their homologs on the X. The more recent theory of Y degeneration by regulatory evolution (Lenormand et al. 2020; Lenormand and Roze 2022) proposes that *cis*-regulatory elements evolve to suppress the expression Y-linked genes, which allows Y-linked mutations to accumulate and, in turn, favours further suppression, and eventually silencing, of these genes. Divergence in the proportions of sex-biased genes carried by an X and Y pair is potentially influenced by several factors (Vicoso and Charlesworth 2006), including chromosomal differences in the rates at which male- and female-biased genes evolutionarily accumulate through gene duplication (Wu and Xu 2003; Gallach and Betrán 2011; Connallon and Clark 2011), translocations or centric fusions between sex chromosomes and autosomes (Charlesworth and Charlesworth 1980; Pennell et al. 2015), *de novo* gene formation (Begun et al. 2007), and gene regulatory divergence leading to sex-biased gene expression (Rice 1984; Connallon and Clark 2010). Regulatory constraints associated with dosage compensation and meiotic X inactivation—as observed in some mature sex chromosome systems—might further contribute by making the X a transcriptionally inhospitable environment for male-biased genes (Vibranovski et al. 2009; Bachtrog et al. 2010; Meisel et al. 2022).

Notably, while each of the scenarios mentioned above can influence specific features of sex chromosome evolution, several must act concurrently to explain the overall patterns of gene content observed on the X and Y. Here, we argue that an extended version of Muller’s classic “sheltering” hypothesis (Muller 1914, 1918; Muller and Painter 1932) might also contribute to the evolutionary patterns of both gene loss and sex-biased gene content on sex chromosome. Since Muller’s original hypothesis was disregarded decades ago, we begin by presenting a brief history of Muller’s hypothesis along with the major arguments that led to its rejection. It should be noted that Muller’s sheltering scenario is unrelated to the recent discussion about the role of “sheltering” in recombination suppression between the X and Y chromosome (Jay et al. 2024; Charlesworth and Olito 2024). We then present an extended mathematical model that relaxes restrictive assumptions of earlier sheltering models and shows that sheltering can contribute to the evolution of X- and Y-linked gene losses, along with enrichments of female- and male-biased genes on the X and Y chromosome, respectively (our models also apply to Z and W chromosomes once sex labels are switched). Our model clarifies the conditions under which sheltering should be an important factor in sex chromosome evolution. Finally, we review contemporary data on gene expression, sex-linked gene content, and the fitness effects of loss-of-function (LOF) mutations, in a broader discussion of the potential contribution of sheltering to sex chromosome evolution.

### The rise and fall of Muller’s sheltering hypothesis

In 1914, H. J. Muller (co-crediting fly-room colleague Alfred Sturtevant) proposed the “sheltering hypothesis” for Y chromosome degeneration (Muller 1914; Muller 1918; Muller and Painter 1932; Fisher 1935; Nei 1970). Muller argued that an ancestral Y chromosome— which contains the same set of genes but lacks recombination with the ancestral X—should degenerate over time through the accumulation of recessive Y-linked mutations that result in a loss-of-function (LOF) for the gene (such mutations, in contemporary language, include nonsense mutations, frameshifts, and whole-gene deletions). If, as Muller predicted, Y-linked LOF mutations are invariably heterozygous with functional X-linked copies of the same genes, then the fitness effects of these Y-linked variants will be sheltered from selection. Such mutations can, therefore, become fixed on the Y chromosome, which decays in the number of functional genes it carries without consequences for male fitness.

Muller’s hypothesis for Y chromosome degeneration nevertheless fell out of favour during the latter half of the 20^th^ Century, and when mentioned today (which is rare), is usually summarily dismissed as little more than a historical footnote (*e.g.*, Charlesworth 1978, 1991; Rice 1996; Orr and Kim 1998; Abbott et al. 2017; Vicoso 2019). This was due to three main arguments. The first, and most convincing, was that formal population genetic models contradicted Muller’s intuition about the evolutionary consequences of sheltering. Muller correctly noted that Y-linked mutations can be masked by functional copies of the same genes on the X, yet he erred in assuming that such sheltering effects are sufficient to cause Y-linked gene loss. Using a deterministic model, Fisher (1935) showed that recessive lethal mutations cannot reach high frequencies on the Y chromosome because similar mutations also arise on the X and render Y-linked sheltering effects incomplete: when X- and Y-linked mutation rates are equal, lethal alleles evolve to identical equilibrium frequencies on the X and Y and remain rare on both chromosome types. Extensions of the model to include male-biased mutation rates (Fisher 1935) and inbreeding (Frota-Pessoa and Aratangy 1968) permit lethal alleles to reach higher frequencies on the Y than the X, though these conditions are still not sufficient for lethals to become fixed. Finally, Nei (1970) explored the effect of genetic drift on the fixation rates of Y-linked lethals and found that the population size must be exceptionally small for fixation to be likely, which precludes a role for sheltering in species with large population sizes, of which there are many (e.g., Buffalo 2021). Collectively, these early attempts to formally model sheltering convinced many of the inadequacy of the mechanism (Charlesworth 1978, 1991; Rice 1996; Orr and Kim 1998).

Charlesworth (1978) raised two further arguments against the sheltering hypothesis, which revolved around the empirical evidence for dosage compensation of the X chromosome and the dominance of deleterious mutations. Sheltering effects are strongest when mutations are completely recessive (*i.e.*, when they have dominance coefficients of *h* = 0), whereas incomplete recessivity (dominance coefficients within the range: 0 < *h* < 0.5) leads to selection against heterozygous individuals and reduces the capacity of sheltering to cause Y chromosome degeneration (Nei 1970). Mutation accumulation data available during the 1970s showed that the *average* dominance coefficient of mildly deleterious mutations was within the range 0.1 < *h* < 0.5 (Simmons and Crow 1977; Crow 1993), while estimates of the mean dominance of lethal mutations (as possible proxies for LOF alleles in general) suggested even stronger, but on average incomplete, recessivity (i.e., 0.01 < *h* < 0.05 on average; Simmons and Crow 1977; Crow 1993). Although most of the available mutation data were insufficient for estimating other aspects of the distribution of dominance (e.g., variance, skew, or the proportion of mutations with nearly complete recessivity; Halligan and Keightley 2009; we expand upon the limits of contemporary data on dominance in the Discussion), the (non-zero) estimates of mean dominance nevertheless reinforced the view espoused by the early mathematical models: that sheltering was unlikely to be important in sex chromosome evolution (Charlesworth 1978, 1991; Rice 1996; Orr and Kim 1998).

The final argument is based on the observation that species with highly degenerate Y chromosomes often evolve dosage compensation, which implies that there is a fitness cost to males of gene loss from the Y (Charlesworth 1978, 1991). This argument rules out sheltering as a universal explanation for Y chromosome degeneration in species that have evolved dosage compensation, yet it does not rule out significant contributions of sheltering to sex chromosome evolution alongside other processes (*e.g.*, selective interference: Charlesworth 1978; Bachtrog 2013; regulatory evolution: Lenormand and Roze 2022) that do favour the evolution of dosage compensation. The observation of dosage compensation, likewise, says nothing about the proportion of genes that might incur negligible fitness costs in hemizygous state. Indeed, contemporary examples of species with highly degenerate Y or W chromosomes that have not evolved complete dosage compensation on the X or Z, or where compensation occurs on a gene-by-gene basis (e.g., Mank et al. 2011; Gu and Walters 2017; Zhu et al. 2024), imply that dosage-related fitness costs of Y- or W-linked gene loss might sometimes be negligible.

### An extended model of X and Y chromosome sheltering

#### Preliminary Comments

The aim of our analysis is to relax two restrictive assumptions of prior models, which may have led to an overly pessimistic view of the potential contribution of sheltering to sex chromosome evolution. Firstly, previous models largely focused on the potential for fixation of homozygous lethal mutations, though we now know that most genes in multicellular Eukaryotes are non-essential (i.e., their LOF alleles are not lethal; Rancati et al. 2018). Second, and more importantly, previous models invariably assumed that LOF mutations have identical homozygous fitness costs in each sex. While this assumption is reasonable for lethal mutations (a large proportion of which affect both sexes; Ashburner et al. 2005), it is unlikely to hold for LOF mutations in general. For example, mutations conferring sterility in one sex do not typically cause sterility in the other (Lindsley and Lifschytz 1972), and spontaneous mutations often differentially affect the fitness of each sex (Mallet et al. 2011; Sharp and Agrawal 2013). Moreover, two decades of transcriptomics research shows that large fractions of the genome exhibit sex-biased gene expression (Ellegren and Parsch 2007; Parch and Ellegren 2013; Grath and Parsch 2016) and this sexually dimorphic gene expression is at least somewhat indicative of sex differences in a gene’s functional importance (Connallon and Clark 2011). Genes with sex-limited expression also appear to be common within animal genomes (Perry et al. 2014; Mongue et al. 2021; Bain et al. 2021; Yu et al. 2023; see our Discussion) and must, by definition, have sex-limited fitness effects. It therefore seems plausible for LOF alleles to differentially affect the fitness of each sex, though this has not yet been systematically tested.

While previous sheltering models focused on Y-linked gene loss (Fisher 1935; Frota-Pessoa and Aratangy 1968; Nei 1970), recent theory by Mrnjavac et al. (2023) shows that X-linked deleterious mutations—particularly mutations with fitness costs in males but not females—can be strongly sheltered from selection when the Y carries functional (wild-type) copies of the same genes, potentially leading to decay of male-limited genes from young X chromosomes. Even more recently, Lenormand and Roze (2025) have shown by simulations in small populations (*N* = 10,000) that male-limited genes can degenerate from either the X or the Y chromosome. Each of these results is reminiscent of the strong effects of sheltering that can occur for Y-linked mutations in small populations, where deleterious mutations are unlikely to simultaneously segregate for X- and Y-linked gametologs of individual genes, which is the main factor inhibiting gene loss by sheltering in large populations (cf. Nei 1970; Fisher 1935). What remains to be shown is whether there are selective scenarios that allow for degeneration through sheltering of undifferentiated X and Y chromosomes when X- and Y-linked deleterious alleles co-segregate, and how such a process is shaped by varying population sizes.

We therefore consider the evolutionary accumulation of mutations within homologous regions of an X and Y chromosome pair that do not recombine and initially carry the same set of functional genes (for full details of the model, see Appendix 1-4 of the Supporting Information). We focus on LOF mutations given the widespread empirical observation that LOF alleles are the most likely genetic variants to be strongly recessive (Agrawal and Whitlock 2011; Manna et al. 2012; Balick et al. 2022), which is an essential feature of all sheltering models. Our most important departure from previous models of sheltering (Fisher 1935; Frota-Pessoa and Aratangy 1968; Nei 1970) is that we allow mutations to differentially affect female and male fitness. We later consider how sheltering effects interact with evolutionary models of Y chromosome degeneration that were developed after sheltering theory was rejected (i.e., from the 1970s onwards). These include models of selective interference on nonrecombining Y chromosomes (Charlesworth 1978; Bachtrog 2006, 2013; Charlesworth and Campos 2014), and recent models of Y degeneration by regulatory evolution (Lenormand et al. 2020; Lenormand and Roze 2022).

#### Deterministic evolutionary dynamics of LOF mutations on the X and Y chromosome

We begin by considering a deterministic model of recurrent mutation and selection at single genes and later consider the effects of drift, selective interference, and regulatory evolution on the fixation of LOF alleles. We assume that functional copies of each gene mutate to LOF alleles at rates μ_*f*_ and μ_*m*_ in females and males, respectively, with back-mutation assumed to be small enough to be negligible. Because there are many ways for a single gene to lose function, from single-nucleotide mutations that introduce premature stop codons to deletions or insertions that disrupt the protein or its regulatory sequences, genic LOF mutation rates are thought to be high, with proposed rates on the order of μ ∼10^-4^ to 10^-6^ (Drake et al. 1998; Monroe et al. 2021; Balick et al. 2022). LOF alleles of a given gene reduce fitness by *s_f_* and *s_m_* in female and male homozygotes (0 ≤ *s*_*f*_, *s*_*m*_ ≤ 1) and by *s_f_h* and *s_m_h* in heterozygotes, where *h* is the dominance coefficient (0 < ℎ < 0.5 corresponds to partial, and *h* = 0 to complete, recessivity). We assume throughout that the sex-averaged strength of selection against LOF mutations is much larger than sex-averaged mutation rates (*s*^-^ ≫ μ^-^, where *s*^-^ = (*s*_*f*_ + *s*_*m*_0⁄2 and μ^-^ = (μ_*f*_ + μ_*m*_0⁄2). These conditions prevent gene losses on autosomes, though as we shall see, they permit gene losses on sex chromosomes.

When mutations are recessive and fitness effects are equally strong in each sex (*h* = 0 and *s*_*f*_ = *s*_*m*_), we recapture the results of earlier sheltering models for very large populations (i.e., effectively deterministic; Fisher 1935). In this case, LOF mutations remain rare on both the X and the Y. As noted by Fisher (1935), the mutation-selection equilibria are identical between the X and Y when mutation rates are also equal between the sexes (*i.e.*, *p^^^*_*x*_ = *p^^^*_y_ when μ_*f*_ = μ_*m*_ and *s*_*f*_ = *s*_*m*_, where *p^^^*_*x*_ and *p^^^*_y_ represent the equilibrium LOF allele frequencies on the X and Y, respectively; see Fig. 1). Male-biased mutation rates within the range that is typically observed in animals (*i.e.*, μ_*f*_ ≤ μ_*m*_ ≤ 4μ_*f*_; Connallon et al. 2022) elevate LOF allele frequencies on the Y relative to the X, though such alleles remain rare on both chromosomes.

**Figure 1.**
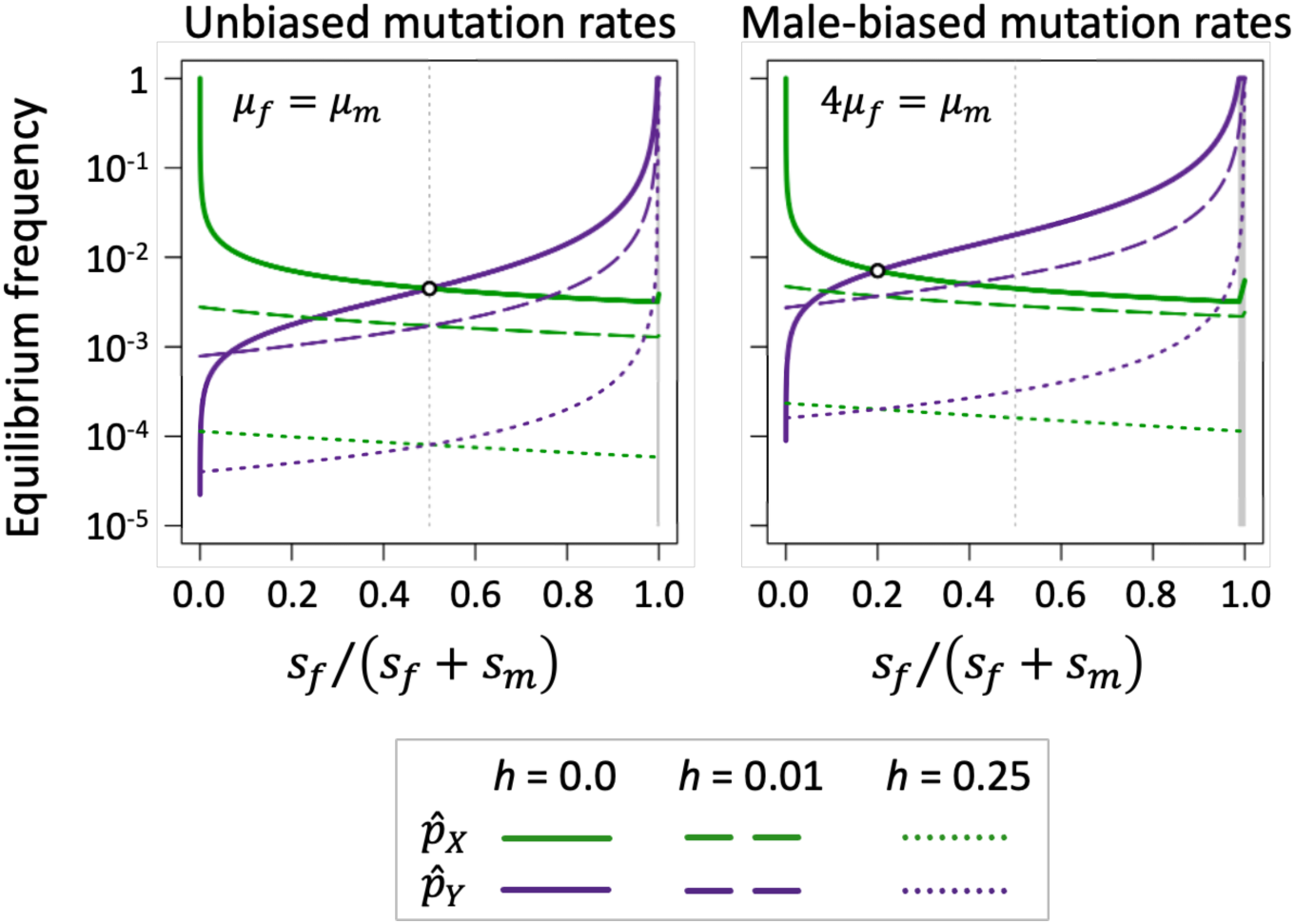
Sexually dimorphic fitness effects of LOF mutations promote sex chromosome gene losses. Each panel shows mutation-selection equilibrium frequencies (in log_10_ scale) for X- and Y-linked genes under different scenarios of dominance for LOF mutations (completely recessive: *h* = 0; partially recessive: *h* = 0.01 and *h* = 0.25). The *x*-axis spans the gradient between male-limited genes (*s_y_* /(*s_y_* + *s_m_*) = 0 when *s_m_* > 0 and *s_y_* = 0), genes that with equal fitness effects in each sex (*s_y_* /(*s_y_* + *s_m_*) = 0.5 when *s_m_* = *s_y_*) and female-limited genes (*s_y_* /(*s_y_* + *s_m_*) = 1 when *s_m_* = 0 and *s_y_* > 0). The shaded grey regions show parameter space favouring deterministic fixation of recessive LOF mutations on the Y chromosome; the parameter region leading to X-linked gene losses is much narrower and includes genes with male-limited functions. Results show cases where *s*^-^ = 0.5 and μ_*f*_ = 10^-5^. Equilibria were determined analytically in the cases where LOF mutations were completely recessive (h = 0), and they were otherwise determined numerically.

Sexually dimorphic fitness costs allow LOF mutations to differentially accumulate on the X or Y (Fig. 1). LOF mutations become enriched on the X chromosome in genes that are disproportionately important for male fitness (*p^^^*_*x*_ > *p^^^*_y_ for genes where *s*_*m*_⁄*s*_*f*_ > μ_*m*_⁄μ_*f*_), while LOF mutations preferentially accumulate on the Y in genes that are disproportionately important for female fitness (*p^^^*_*x*_ < *p^^^*_y_ for genes where *s*_*m*_⁄*s*_*f*_ < μ_*m*_⁄μ_*f*_). Complete fixation of a LOF allele on the X or the Y (*i.e.*, *p^^^*_*x*_ = 1 or *p^^^*_y_ = 1) occurs when sheltering effects are strong and LOF alleles have sufficiently pronounced sexual dimorphism in their homozygous fitness effects. Completely recessive mutations become fixed in X-linked genes with male-limited functions (*i.e.*, *p^^^*_*x*_ = 1 when *h* = 0 and *s*_*f*_ ≤ μ_*f*_). A modest amount of expression in females (*s*_*f*_ > μ_*f*_) or in heterozygotes (*e.g*., *h* = 0.01 in Fig. 1) is sufficient to prevent fixation of X-linked LOF mutations.

Conditions for fixation of Y-linked LOF mutations are much more permissive. Unsurprisingly, female-limited genes (those where *s*_*m*_ = 0) decay from the Y regardless of their fitness effects in females. However, an absence of expression in males is not required for fixation of Y-linked LOF alleles. Indeed, recessive LOF allele fixation occurs when *s*_*m*_ < μ_*m*_A*s*_*f*_⁄μ_*f*_, which corresponds to the grey shaded regions in Fig. 1. Consequently, genes that are essential for females (*s*_*f*_ = 1) can deterministically decay from the Y even if they have moderate fitness costs in males (e.g., fitness effects greater than 1% when μ_*f*_ = 10^-5^and μ_*m*_ = 4μ_*f*_; higher LOF mutation rates further expand the range male fitness costs that are compatible with Y-linked gene loss). Incomplete recessivity (*h* > 0) reduces the scope for Y-linked LOF allele fixation (though gene losses remain possible), whereas male-biased mutation rates expand the scope (Fig. 1).

#### Effects of genetic drift and sheltering on X and Y chromosome gene losses

We next evaluated effects of genetic drift on fixation of recessive LOF alleles using a combination of analytical and simulation approaches (see Appendix 2 of the Supporting Information; incompletely recessive alleles are considered further below). We first used a diffusion approximation to calculate the fixation probabilities for new Y-linked recessive LOF mutations that enter a population in which X-linked genetic variation segregates at mutation-selection-drift balance. Following Nei (1970), we assumed that X-linked variation evolved in the absence of ancestral Y-linked variants, which allows X-linked mutations to reach higher frequencies than they would if the Y were polymorphic. This assumption reduces sheltering effects for Y-linked variants and makes the following results conservative with respect to the fixation probabilities of Y-linked mutations. Secondly, we carried out full stochastic forward simulations in which recurrent mutation, selection, and genetic drift, affect the evolutionary dynamics of LOF alleles at both X- and Y-linked genes.

Genetic drift alters the predictions of our deterministic model in two important ways. First, drift expands the scope for LOF allele fixation on the Y chromosome, owing to its relatively small effective population size (effective population sizes are taken to be 2*N_e_* for autosomes, 1.5*N_e_* for the X, and 0.5*N_e_* for the Y; Hartl and Clark 2007). Y-linked recessive LOF mutations not only fix with high probabilities (i.e., of similar order to neutral mutations) under parameter conditions leading to deterministic degeneration on the Y, but they also fix under parameter conditions which do not favour deterministic fixation (Fig. 2). As in the deterministic model, genes with strongly female-biased functions are consistently lost from the Y. In addition, LOF mutations that substantially reduce fitness of both sexes often fix in cases where the population-scaled LOF mutation rate per gene is small (*i.e.*, *N*_*e*_μ^-^ ≪ 1; Fig. 2), which applies to small genes (where μ^-^ is small) and lineages with small effective population sizes.

**Figure 2.**
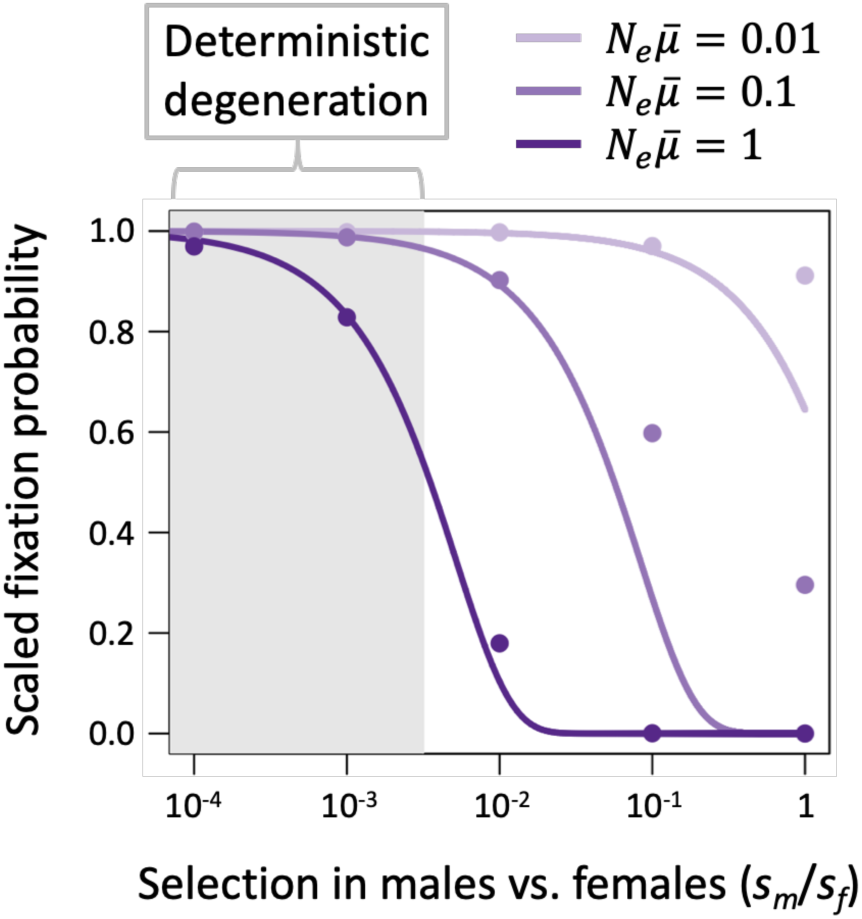
Fixation probabilities of Y-linked LOF mutations depend on interactions between drift and sex differences in selection. The curves show analytical predictions for unique Y-linked mutations entering a population at mutation-selection-drift equilibrium for X, with no ancestral genetic variation on the Y (based on eq. (2) with *f*_0_ = 1). The circles show proportions of Y-linked LOF mutations that fix in replicate computer simulations that incorporate selection and drift on X and Y chromosomes, and recurrent mutation on the X. Each datapoint is based on 10^7^-10^9^ replicate simulations ending in fixation or loss of the Y-linked variant. All fixation probabilities are scaled relative to those of neutral mutations. The shaded region represents the parameter space leading to deterministic fixation (which corresponds to the condition *s*_*m*_ < μ_*m*_A*s*_*f*_⁄μ_*f*_). Results are shown for a LOF mutation rate of μ^-^ = μ_*f*_ = μ_*m*_ = 10^-5^and three population sizes (*N* = *N*_*e*_ = 10^5^, 10^4^, and 10^3^). LOF alleles are completely recessive (*h* = 0) and genes are essential for females (*s_y_* = 1).

Second, genetic drift affects the predictability with which male-limited genes are lost from the X versus the Y chromosome (Fig. 3). Male-limited genes with intermediate-to-large population-scaled mutation rates (*e.g.*, *N*_*e*_μ^-^ > 0.2) and recessive LOF alleles (*h* = 0), are reliably lost from the X and retained on the Y, consistent with deterministic predictions. By contrast, small population-scaled mutation rates lead to a mix of male-limited gene losses from the X and Y, which reflects the strong sheltering effects that arise on both chromosomes in cases where deleterious mutations rarely segregate simultaneously on the X and Y (Fig. 3A). As the population-scaled LOF mutation rate approaches zero (*N*_*e*_μ^-^ → 0), the proportion of male-limited genes lost from the X versus the Y approaches the limit *f*_*x*_ = (2 + μ_*m*_⁄μ_*f*_0/(2 + 4μ_*m*_⁄μ_*f*_0, in which at least half of all gene losses are Y-linked (*f*_*x*_ = 0.5 when μ_*m*_ = μ_*f*_; *f*_*x*_ < 0.5 when μ_*m*_ > μ_*f*_; Fig. 3A). The evolutionary rates at which sex-limited genes become lost from X and Y chromosomes are also predictable and rapid relative to the age of many sex chromosome systems (Fig. 3B Fig. S1 of the Supporting Information). For male-limited genes on the X, the average frequency of LOF mutations *t* generations following suppression of recombination between the X and Y is approximately *p*_*x,t*_ = 1 − *e*^-2μ*ft*⁄3^, and for female-limited genes on the Y, the average frequency of LOF mutations is *p*_F,*t*_ = 1 − *e*^-μ*mt*^ (see Appendix 2 of the Supporting Information).

**Figure 3.**
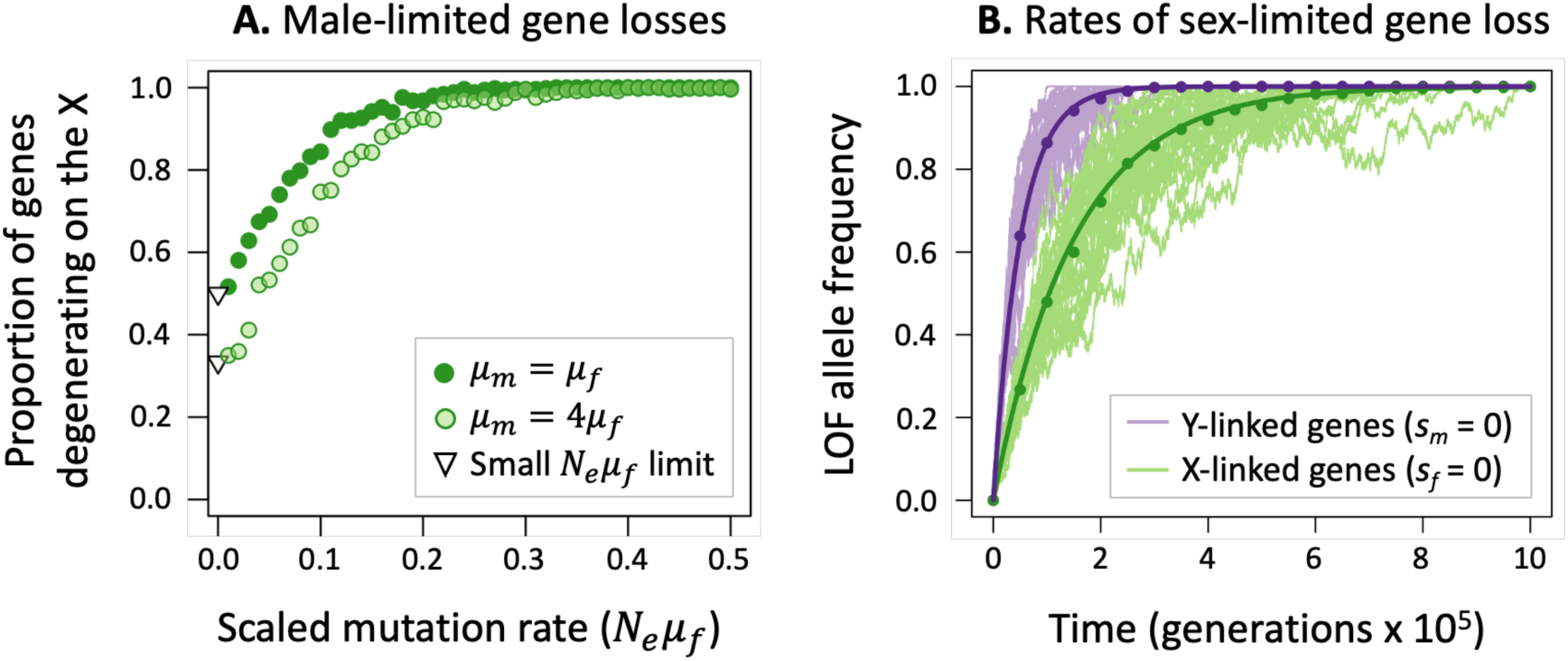
Genetic drift affects the predictability of X- and Y-linked gene loss. Panel A: Genes with male-limited functions eventually degenerate from either the Y or the X chromosome, owing to fixation of recessive LOF alleles. In each of 500 replicate simulations, LOF mutations were initially absent from the X and Y; both chromosomes were permitted to evolve under recurrent mutation, selection and drift, until a LOF allele was fixed on the X or the Y (with *h* = 0, μ_*f*_ = 10^-4^, *s_m_* = 0.3, *s_y_* = 0, *N_e_* ranging from 10^2^ to 10^4^). Circles shows the proportion of male-limited genes degenerating from the X but retained on the Y; the remaining proportion degenerate from the Y and retained on the X. Triangles show the limit in which the population-scaled mutation rate approaches zero. **Panel B:** Recessive LOF mutations eventually fix in sex-limited genes, with drift causing variation in evolutionary trajectories of LOF alleles over time. Results show examples of degeneration in a large population (*N_e_* = 10^5^) with parameters μ_*f*_ = 10^-5^, μ_*m*_ = 2μ_*f*_, *s_y_* = 1 and *s_m_* = 0 for Y-linked genes, and *s_m_* = 1 and *s_y_* = 0 for X-linked genes. 30 simulation runs were carried out for each chromosome, with individual trajectories (thin, pale lines) scattered about the analytical predictions (bold curves) and circles denoting mean LOF frequencies across the set of simulated trajectories.

These predictions apply when LOF alleles are *effectively* recessive (i.e., effectively neutral in heterozygous but not homozygous state: *N*_*e*_*s* ≫ 1 and *N*_*e*_*s*ℎ < 1). Higher degrees of dominance render LOF mutations unlikely to fix, particularly when they are X-linked, though as we illustrate below, sheltering effects remain permissive on the Y in concert with other processes contributing to Y chromosome degeneration.

#### Interactions between sheltering and other processes of Y chromosome gene loss

The decline of the sheltering hypothesis coincided with the rise of models of Y chromosome decay by selective interference (due to hitchhiking effects of selective sweeps and background selection; see Bachtrog 2006, 2013; Charlesworth and Campos 2014) and, more recently, the theory of Y degeneration by regulatory evolution (Lenormand et al. 2020; Lenormand and Roze 2022). Consequently, there is currently no theory evaluating the interactions between sheltering and these other scenarios of Y-linked gene loss. We therefore sought to quantify how these more recent scenarios modulate the fixation of LOF alleles on the Y. For clarity, we separately evaluate interactions between sheltering and each contemporary mechanism of Y degeneration (with derivations provided in Appendix 3 of the Supporting Information), though we note that all can occur simultaneously in nature (see Bachtrog 2008; Lenormand et al. 2020; Lenormand and Roze 2022).

The interaction between sheltering and degeneration by regulatory evolution (DRE) is straightforward. The DRE model predicts that *cis*-regulatory regions will evolve to over-express X-linked and under-express Y-linked gametologs, rendering Y-linked mutations more recessive than they would otherwise be on the X or autosomes (Lenormand et al. 2020; Lenormand and Roze 2022). This effect lowers the effective dominance of deleterious mutations, making fixation more likely. LOF mutations are already expected to have strongly recessive effects (see the discussion below). The extent to which *cis*-regulatory elements evolve to decrease Y-relative to X-linked gene expression should further dampen any harmful effects of Y-linked LOF mutations in males and, thus, increase their probabilities of fixation.

Selective interference on the Y should also enhance the fixation probabilities of LOF alleles. In cases where a beneficial mutation arises on a Y chromosome that carries one or more LOF alleles, a selective sweep of the beneficial variant will carry linked LOF alleles to fixation, resulting in Y-linked gene loss. Assuming that LOF alleles are recessive, fitness effects of new beneficial mutations are exponentially distributed with a mean of *s*^-^_*b*_, and LOF alleles of a gene initially segregate at mutation-selection equilibrium (*p^^^*_y_ = μ_*m*_⁄(*s*_*m*_*p^^^*_*x*_) and *p^^^*_*x*_ = Aμ_*f*_⁄*s*_*f*_ on the Y and X, respectively), then the probability that the beneficial mutation arises in association with a LOF allele at a single gene and then carries it fixation is:

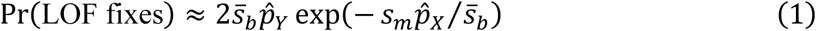

(see Appendix 3 of the Supporting Information). Numerical evaluation of eq. (1) and stochastic simulations (Fig. 4A) show that hitchhiking events substantially elevate fixation probabilities of Y-linked LOF alleles relative to those of neutral mutations (Fig. 4A), which expands the parameter space of selection over which LOF allele fixation is likely. The effect of hitchhiking is most pronounced for genes that are more important for female than male fitness (*i.e.*, genes where *s_m_*/*s_f_* < 1), though fixation probabilities for LOF alleles remain high in cases where the gene is essential for both sexes (see *s_m_*/*s_f_* = 1 in Fig. 4A, which corresponds to lethal mutations). These hitchhiking predictions extend to cases where multiple loci segregate for LOF alleles (Fig. 4B; see Appendix 3 of the Supporting Information).

**Figure 4.**
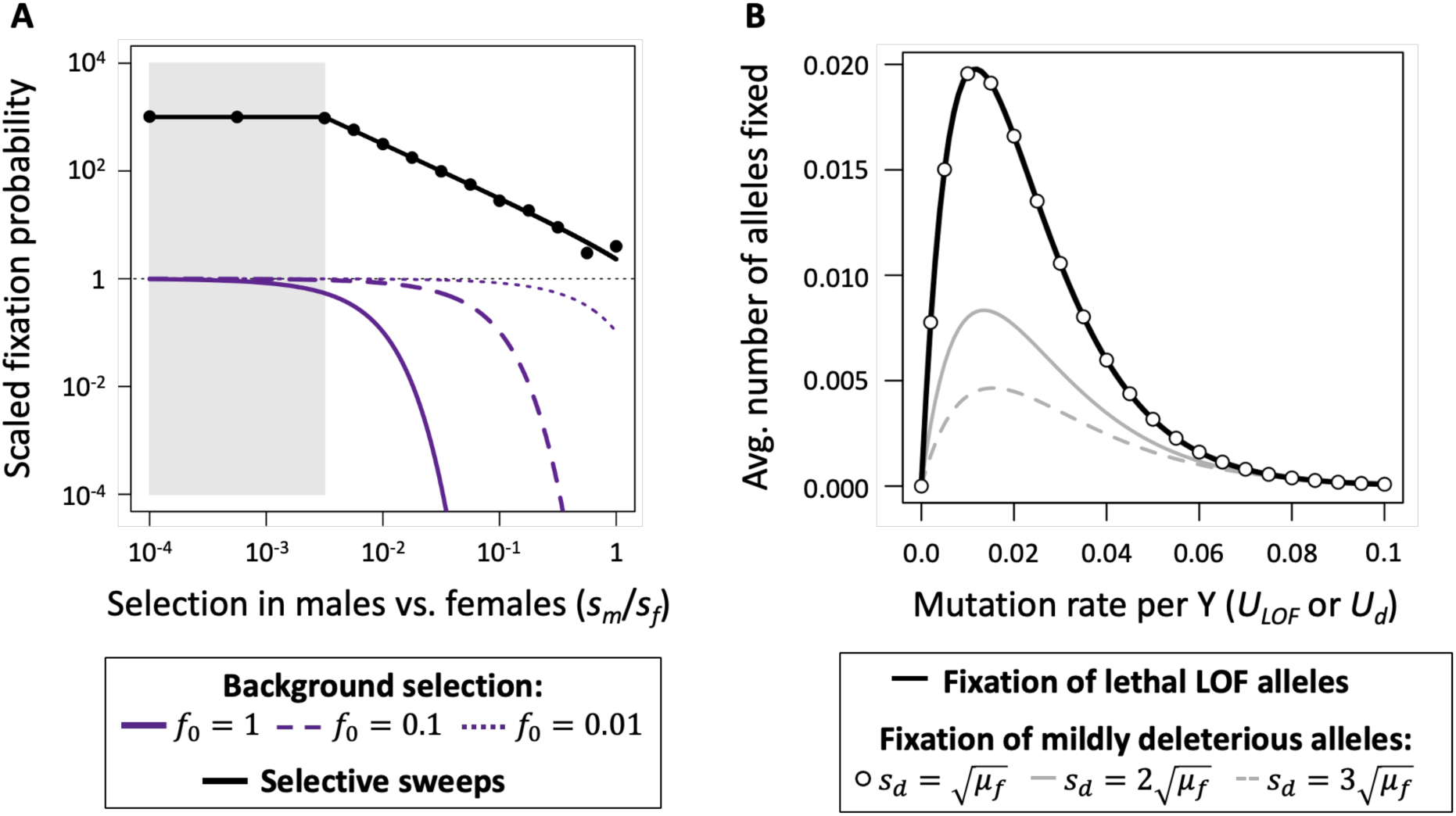
Conditions for recessive Y-linked LOF allele fixation through hitchhiking and background selection. Panel A: The grey shaded region shows the parameter space for deterministic fixation of recessive LOF alleles (*s*_*m*_ < μ_*m*_A*s*_*f*_⁄μ_*f*_). The solid black line (eq. (1)) shows the probability that a LOF allele at a single Y-linked gene hitchhikes to fixation with a beneficial mutation. Circles show the proportion of 10^5^ simulated beneficial mutations that sweep to fixation with a LOF allele. Purple curves (eq. (2)) show the fixation probabilities a new Y-linked LOF mutations under three levels of background selection *f*_0_, which represents proportion of Y chromosomes that are free of deleterious mutations. Fixation probabilities in panel A are scaled relative to those of neutral mutations (*i.e.*, 2/*N* for Y-linked mutations, shown by the horizontal broken black line). **Panel B:** effects of hitchhiking when many deleterious variants segregate simultaneously on the Y chromosome. The black curve shows cases where recessive lethal alleles segregate at mutation-selection balance and potentially hitchhike with a beneficial mutation. The remaining results show cases of hitchhiking of mildly deleterious mutations with heterozygous effects of *s_d_*. Curves are based on equations presented in Appendix 3 of the Supporting Information. Results use the parameters μ^-^ = μ_*f*_ = μ_*m*_ = 10^-5^, *N* = *N*_*e*_ = 10^5^, *s_y_* = 1, and *s*^-^_*b*_ = 0.01.

Background selection similarly enhances fixation of recessive LOF alleles. We suppose that LOF alleles are initially absent from the Y and segregating at mutation-selection-drift balance on the X (as in the analytical results of Fig. 2), and we let *f*_0_ represent the proportion of Y chromosomes that is free of background deleterious genetic variation (*i.e.*, semidominant deleterious variants maintained at mutation-selection equilibrium). Following the general approach of Charlesworth (1994), the probability that new, recessive LOF alleles become fixed on the Y is:

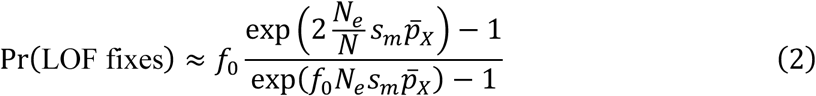

(Appendix 3, Supporting Information), where *p*^-^ = Γ O1.5*N* μ + <u>^1^</u>P OA*N s* Γ(1.5*N* μ)P is the expected frequency of the LOF allele on the X, Γ(*x*) refers to the gamma function, *N_e_* is the effective population size, and *N* is the census size. Numerical evaluation of eq. (2) shows that background selection on the Y, which reduces *f*_0_, can greatly expand the conditions under which LOF alleles can fix (Fig. 4A).

As selective interference enhances the fixation rates of all classes of deleterious mutation, the question is how the fixation probabilities of recessive LOF alleles compare to those of mildly deleterious Y-linked mutations? Fixation probabilities of mildly deleterious Y-linked mutations depend on their heterozygous fitness effect on males (which we can define as *s_d_*), whereas the probabilities for recessive LOF mutations depend on their homozygous costs in males (*s_m_*) weighted by the frequency of LOF alleles for the X-linked copy of the gene (*p*_*x*_). Recessive LOF alleles fix more readily than mildly deleterious mutations whenever *s*_*m*_*p*_*x*_ < *s*_*d*_. For the extreme case of genes that are essential for both sexes (*s_f_* = *s_m_* = 1) and the population is at mutation-selection equilibrium (at *p^^^*_*x*_ and *p^^^*_y_), selective sweeps will more readily fix LOF alleles when *s*_*d*_ > Aμ_*f*_, where μ_*f*_is the genic LOF mutation rate (Fig. 4B; Appendix 3, Supporting Information). Conditions for LOF allele fixation are even more permissive when they are more harmful for females than males (*s_f_* > *s_m_*) and/or drift causes the frequency X-linked LOF alleles to drop below their mutation-selection equilibria (*p*_*x*_ < *p^^^*_*x*_). Comparable results apply under background selection (recessive LOF mutations fix more readily than mildly deleterious mutations when *s*_*d*_ > *s*_*m*_*p*^-^_*x*_).

While incomplete recessivity (*h* > 0) lowers the probability of LOF allele fixation, fixation probabilities can remain substantial in large populations in the presence of selective interference. For example, in cases where LOF alleles are not segregating on the X, Y-linked LOF mutations have moderate fixation rates (of at least one-tenth the neutral fixation probability) if their dominance coefficients fall below the threshold *h*_*crit*_ = 3.6⁄(*N*_*e*_*f*_0_*s*_*m*_) (see Appendix 4 of the Supporting Information), where *f*_0_ again captures the strength of background selection on the Y. Conditions for fixation remain permissive when *f*_0_*s*_*m*_ is small, which is plausible for genes that are non-essential in males (e.g., *s*_*m*_ ≪ 1) and selective interference is strong cross the Y (e.g., *f*_0_ ≪ 1; see Fig. 5A). Probabilities of Y-linked gene loss are dampened when LOF mutations also segregate on the X, yet the degree of dampening remains sensitive to the strength of selection in females (Fig. 5B; see Appendix 4 of the Supporting Information). Frequencies of X-linked LOF alleles decrease as the strength of selection in females increases, which increases sheltering of LOF alleles of their gametologs on the Y. This enhancement of Y-linked sheltering through selection on females is relevant for Y-linked alleles with dominance coefficients of *h* < 0.03 (Fig. 5B), which should apply to a large fraction of LOF mutations (see Agrawal and Whitlock 2011; Manna et al. 2012; Balick et al. 2022).

**Figure 5.**
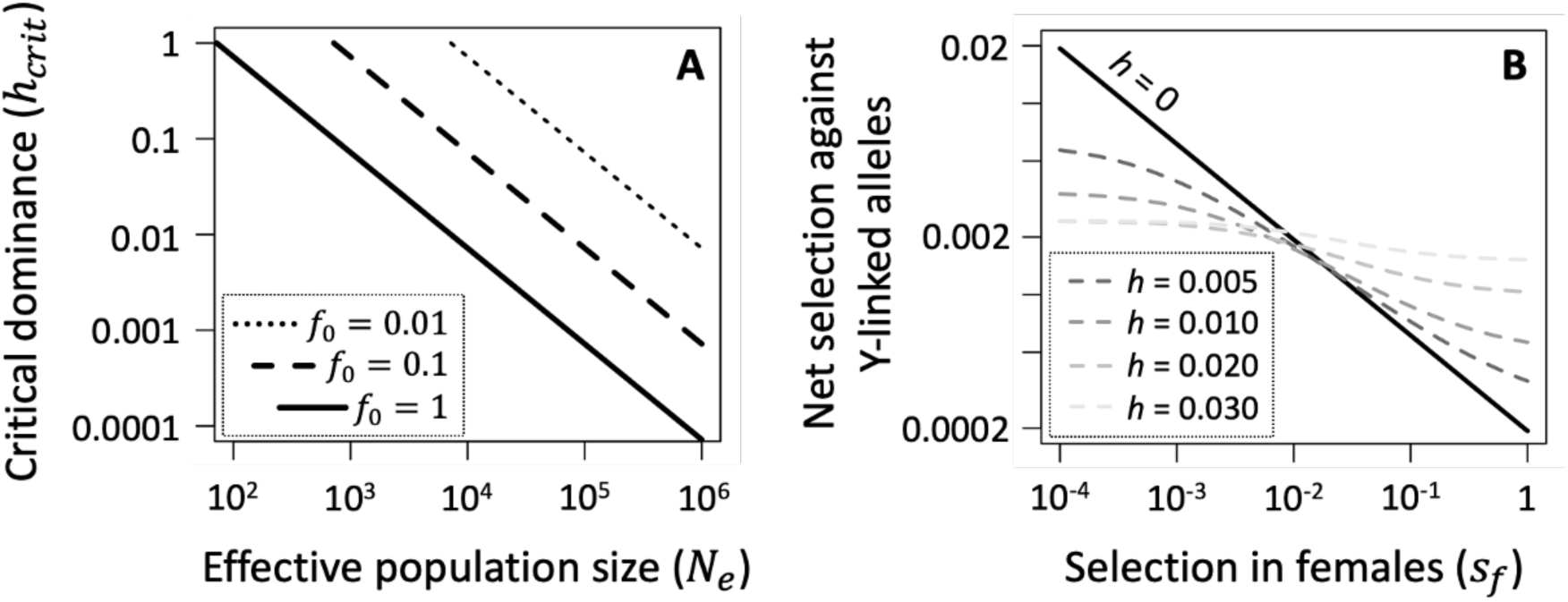
Effects of sheltering on selection against partially recessive Y-linked LOF mutations. Panel A: Dominance thresholds (*h_crit_*) below which LOF alleles are relatively likely to fix. Lines show values of *h_crit_* for Y-linked LOF mutations with homozygous fitness costs of *s_m_* = 0.1, under three degrees of background selection (*f*_0_ = 0 corresponds to no background selection). Results for panel A assume that X-linked genes do not segregate for LOF alleles. Mutations with dominance coefficients below the threshold (*h* < *h_crit_*) have fixation probabilities exceeding 10% of the fixation probability of neutral mutations. **Panel B:** The strength of purifying selection against Y-linked mutations when X-linked LOF alleles segregate at mutation-selection balance. The net strength of selection against a Y-linked mutation is *s*_*m*_\**h* + *p^^^*_*x*_(1 − 2*h*)1, where *p^^^*_*x*_ is the X-linked mutation-selection equilibrium (Appendix 4, Supporting Information). Panel B results show cases where *s*_*m*_ = 0.1 and μ_*f*_ = μ_*m*_ = 10^-5^.

## Discussion

Our model shows that sheltering of LOF mutations promotes the evolutionary decay of certain types of genes from X and Y chromosomes, resulting in enrichments of female-biased genes on the X (owing to a deficit of male-limited genes) and male-biased genes on the Y (owing to a deficit of genes that are more important for female than male fitness). These consequences of sheltering should occur when three conditions are met. First, recombination must be suppressed between homologous regions of the X and Y, as commonly observed in mature sex chromosome systems (with notable exceptions; Perrin 2009; Charlesworth 2021). Second, loss-of-function (LOF) mutations must often be recessive or nearly so—a scenario that, as discussed in detail below, is compatible with current data on dominance and is particularly likely for Y-linked variants (Lenormand et al. 2020), though this issue requires further study. Third, LOF mutations must often exhibit strong sex-biased fitness effects. LOF mutations must be male-limited and completely recessive to reliably fix on the X chromosome. Conditions for fixation are much more permissive for Y-linked genes, including scenarios where LOF alleles are incompletely recessive and X and Y gametologs are expressed by both sexes.

The evolution of separate sexes and sexual dimorphism predates the origin of animal sex chromosome systems, and genes with sex-specific functions were probably prevalent on ancestral autosomes that became sex chromosomes. Multiple lines of evidence suggest that large-effect mutations, including LOF alleles, often have strongly sex-biased fitness effects. In addition to observations that sterility alleles typically have sex-specific effects (Lindsley and Lifschytz 1972; Ashburner et al. 2005) and whole-gene knockouts often have sex-biased or sex-limited phenotypic effects (van der Bijl W, Mank JE. 2021), the pervasiveness of sex-biased gene expression implies that many genes might be sexually dimorphic in their functional importance (Grath and Parsch 2016). While relatively few studies report estimates of the proportion of the genome that is sex-limited in expression, for which we can safely assume that LOF alleles are only costly for one sex, those studies that have indicate substantial proportions of sex-limited genes (*e.g.*, roughly 2-5% of the genome, with male-limited genes often more prevalent than female-limited genes; Perry et al. 2014; Mongue et al. 2021; Bain et al. 2021; Yu et al. 2023). However, for most genes, which are expressed by both sexes, we simply do not have enough data on the fitness effects of LOF alleles to directly evaluate the extent of sexual dimorphism in the fitness costs of LOF alleles. We hope our results will encourage further study of this issue.

To gain a clearer picture of the prevalence of strongly sex-biased and sex-limited genes in animal genomes, we searched for high-quality gene expression studies, prioritizing those using whole-body samples (which should be more conclusive, given our purpose, than studies focusing on a small number of tissues) and when possible multiple life stages. Although our search was not exhaustive (a future systematic survey and analysis would be useful), it includes species spanning several animal phyla, including Arthropoda, Chordata, Platyhelminthes, Nematoda, and Tardigrada (details of the search and analysis can be found in the Supporting Information). We estimated proportions of genes expressed at different degrees of sex-bias, including sex-limited expression, as summarized in Fig. 6. The analysis shows extensive variation among taxa in the degree of sex-biased expression throughout the genome, with several species having large fractions of genes with 5- to 10-fold or higher expression in one sex relative to the other, and other species showing minimal sex-biased expression. There was a clear elevation in the number of strongly male-biased and male-limited genes relative to the number of strongly female-biased and female-limited genes, which hints at a substantial opportunity for the decay of X-linked genes through sheltering effects. Our survey suggests that the processes that we have outlined here may vary in importance among species.

**Figure 6.**
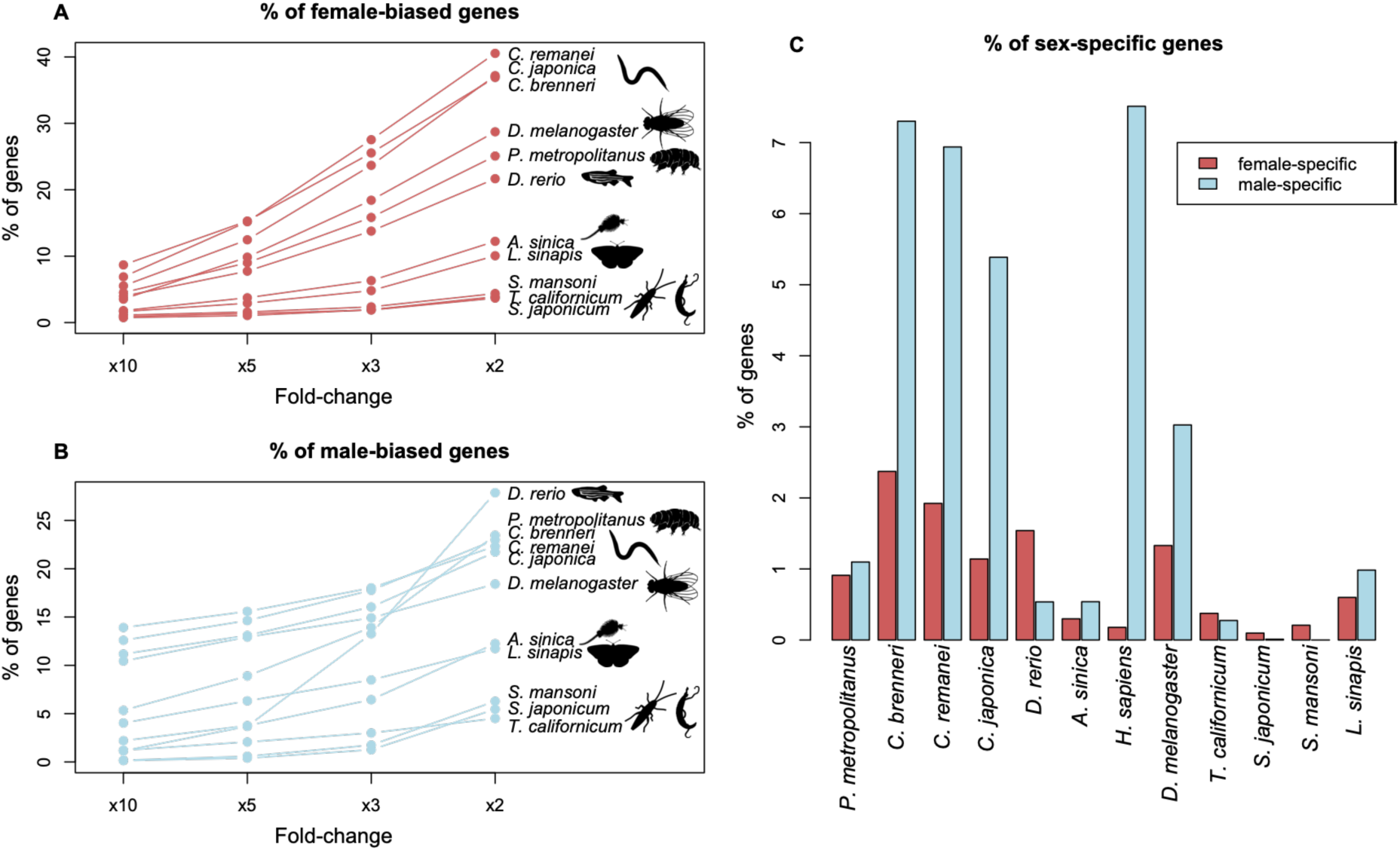
A survey of sex-biased and sex-limited gene expression in animals. Panel A: Percentage of female-biased genes in different species, using various thresholds. **Panel B:** Percentage of male-biased genes at various thresholds. **Panel C:** Percentage of genes with female- and male-specific expression (i.e., >99% of expression is in one of the sexes). Expression data were derived from Thomas et al. (2012), Lonsdale et al. (2013), Leal et al. (2018), Djordjevic et al. (2022), King and Zenker (2023), Elkrewi et al. (2021, 2023), and Sugiura et al. (2024).

### Data on dominance and the potential for strong sheltering of LOF mutations

Data from *Drosophila* and yeast suggest that mean dominance is *h*^U^≈ 0.25 for mildly deleterious mutations (Manna et al. 2011; Charlesworth 2015) and within the range of 0.01 < *h*^U^ < 0.05 for lethal and LOF mutations (Simmons and Crow 1977; Crow 1993; Agrawal and Whitlock 2012; Manna et al. 2012). Nevertheless, estimates of mean dominance tell us little about the broader distribution of dominance, which is unknown (Halligan and Keightley 2009), and leaves open the possibility that a meaningful fraction of LOF mutations might be recessive enough for sheltering to be important in sex chromosome evolution. As Nei (1970) points out in his classic study of sheltering of Y-linked lethals:

> This, of course, does not mean that there are no completely recessive or overdominant lethals. On the contrary, it seems that there are lethals with varying degrees of dominance from slight overdominance to a rather high degree of partial dominance. If this is the case, those genes which are overdominant or completely recessive would be fixed in the population rather quickly but the others would be fixed only slowly, depending on the degree of dominance and population size.

A few studies have reported estimates of higher moments of the distribution of dominance for lethal mutations (*e.g.*, the variance), and these provide information about the types of distributions that might be compatible with data on dominance. Yoshikawa and Mukai (1970) reported point estimates of the mean and variance of dominance for lethal mutations accumulated on 2^nd^ chromosomes of *D. melanogaster* (*h*^U^ = 0.027 and σ^2^ = 0.0027, respectively, which we note are rough estimates). As a thought exercise, if we were to assume that values of *h* for lethal mutations were gamma or a beta distributed (which constrains *h* to be positive), then these point estimates for *h*^U^ and σ^2^ imply a right-skewed distribution with a high proportion of mutations that are nearly recessive (∼60% with 0 < *h* < 0.01). This interpretation aligns with other *Drosophila* data showing that mean dominance of *segregating* lethal alleles is significantly lower than that of new lethal mutations (e.g., Crow 1991, 1993), which implies substantial variation in the dominance coefficients of new lethal mutations.

But given that most genes are not essential (Rancati et al. 2018), and lethal mutations are not necessarily loss-of-function mutations (Marion and Noor 2023), the dominance of lethal mutations may not be representative of LOF alleles in general. More recent, targeted deletion datasets permit direct estimates of the fitness effects of individual loss-of-function alleles.

Using yeast data for non-essential genes, Agrawal and Whitlock (2011) estimated the mean, variance, and skew of the distribution of dominance for whole-gene deletions and found that deletions with large homozygous effects (whose fitness effects can be reliably estimated; see Manna et al. 2012) exhibited a right skewed distribution of dominance with a mean close to but greater than zero. As with lethals, these data are consistent with a high proportion of deletions being near completely recessive, though the direct estimates of fitness from which these inferences are based lack the precision required to rule out evolutionarily meaningful heterozygous fitness effects (i.e., effects of order *sh* >> 1/*N_e_* cannot be ruled out; see Gallet et al. 2012).

Finally, modern population genomics has provided new opportunities for evaluating the fitness effects of deleterious mutations (Eyre-Walker and Keightly 2007), including protein-truncating variants (one type of loss-of-function, or LOF, allele). While assessing the dominance of LOF mutations remains a challenge (see Fuller et al. 2019), analyses of the frequencies of LOF alleles in humans suggest that near complete recessivity is plausible for a large fraction of genes (see Simons and Sella 2016; Balick et al. 2022), though other factors have also been invoked to explain the often high frequencies of human LOF alleles associated with severe homozygous effects (e.g., ascertainment bias: Amorim et al. 2017; overdominant fitness effects: Marion and Noor 2023). A survey of the gnomAD dataset from humans (Karczewski et al. 2020) further shows that accumulation of autosomal LOF mutations is less constrained in genes exhibiting strong sex biased expression than genes with similar expression in each sex (see Fig. S2 of the Supporting Information), which implies a relatively small *N_e_sh* for sex-biased genes that should, if anything, promote X- or Y-linked gene losses by sheltering. Finally, we note that all estimates of dominance apply to autosomal or X-linked loci.

As already mentioned, regulatory processes that suppress expression of Y-linked genes (Lenormand et al. 2020; Lenormand and Roze 2022) should, if anything, cause LOF mutations to be more strongly recessive on the Y than they would be on the X or autosomes, which makes sheltering effects on Y degeneration more plausible than implied by available data on dominance. Altogether, the current data on dominance simply cannot exclude the potential for strong sheltering effects across a meaningful, and potentially large, fraction of LOF mutations.

### Potential contributions of sheltering to sex chromosome evolution

Our model is broadly consistent with several features of sex chromosome gene content evolution. Firstly, sheltering is predicted to contribute to the evolution of Y-linked gene losses and masculinization of the Y chromosome. Deterministic fixation of strongly recessive LOF alleles is predicted in genes that are much more important for females than males (Figs. 1, 5B), with genetic drift and selective interference amplifying the conditions for Y-linked gene loss (Figs. 2-5). Selective interference and regulatory evolution are also expected to cause Y chromosome degeneration (Bachtrog 2006, 2013; Charlesworth and Campos 2014; Lenormand et al. 2020; Lenormand and Rose 2022), though sheltering could contribute alongside these processes to Y-linked gene losses, as effects of all three scenarios are reinforcing.

Secondly, our model predicts the decay of X-linked genes with male-limited functions whose LOF alleles are effectively recessive. For completely recessive LOF alleles, or alleles that are effectively recessive due to *cis*-modifying effects on allelic expression (see Lenormand and Roze 2025), the decay of male-limited genes from the X should be most pronounced in genes or lineages with high population-scaled LOF mutation rates (*N*_*e*_μ^-^ ≫ 0), and otherwise male-limited genes are more likely to be lost from the Y than the X. These predictions support Neuhaus’s (1939) intuition that X-linked male-limited genes might be prone to decay, and they build upon the model of Mrnjavac et al. (2023), which showed that male-limited deleterious mutations are prone to accumulation on X chromosomes paired with an undifferentiated Y. Consistent with our model, X chromosomes often display strong deficits of strongly male-biased genes, particularly so in insects like *Drosophila*, which have historically large effective population sizes, but less so in mammals, where *N_e_* is considerably smaller (Vicoso and Charlesworth 2006). Recent studies of neo-X chromosomes in multiple *Drosophila* species show accelerated pseudogenization rates for genes important for males (these genes remain functional on the neo-Y; Nozawa et al. 2016, 2021). Evidence in vertebrates for X-linked gene losses is sparse, with one exception. The gene *PRSSLY* was present on the ancestral X and Y of mammals and is currently the only gene known to have been lost from the mammalian X and retained on the Y (Hughes et al. 2022). *PRSSLY* has testis-specific expression and is exceptionally long, which implies a large target for LOF mutations. As such, *N*_*e*_μ^-^ may be large for *PRSSLY*, though small for most other mammalian genes, which our model predicts should enhance gene loss on the X.

Thirdly, our model predicts that gene loss can be rapid enough for substantial gene losses in even relatively young sex chromosome systems (reviewed in Charlesworth 2021), including lineages with large population sizes. For example, in populations that are large enough to evolve deterministically, and assuming a genic LOF mutation rate of μ = 10^-5^ per sex, our model predicts that recessive LOF alleles in male-limited genes will reach frequencies of 90% on the X chromosome within 350,000 generations. Using the estimate of 15 generations per year in *Drosophila melanogaster* (Turelli and Hoffmann 1995; Pool 2015), this interval equates to ∼23,000 years, consistent with the rapid gene losses observed on the young neo-X chromosomes of *Drosophila* species (*e.g.*, autosome to sex chromosome fusions occurred at ∼1.1 mya, ∼0.5 mya, and 0.25 mya in *D. miranda*, *D. americana*, and *D. albomicans*, respectively; Nozawa 2021). Deterministic loss of sex-limited genes is even faster on the Y.

In smaller populations that are decidedly non-deterministic and where LOF mutations rarely co-segregate for both the X- and Y-linked gametologs of a gene, substitution rates of LOF mutations can remain relatively high when their fitness effects are incompletely recessive or expressed by both sexes. For example, X-linked LOF mutations in male-limited genes have substitution rates of at least one-tenth the neutral rate when *h* < 3.6⁄(*N*_*e*_*s*_*m*_). Y-linked substitution rates are more permissive, surpassing one-tenth the neutral rate when *h* < 3.6⁄(*N*_*e*_*s*_*m*_*f*_0_), which includes effects of background selection. These rates will of course decline when LOF alleles simultaneously segregate for both the X and Y, yet in such cases, substitution rates of LOF alleles should remain the highest among Y-linked genes whose X-linked gametologs are under strong purifying selection in females. Selection through females promotes masking of Y-linked LOF mutations in males, leading to a more rapid decay of Y-linked genes whose ancestral functions were particularly important for females. This prediction should be testable in taxa with data on the genes initially residing on sex chromosomes and the sex-specific functions of those genes.

## Conclusion

No single process explains all features of sex chromosome evolution. Rather, the key question is who the main players are and how each contributes to empirical patterns of X- and Y-linked gene losses, gains, and regulation. For decades, Muller’s sheltering hypothesis has been dismissed as a potential contributor to sex chromosome evolution. We have shown that an extension of Muller’s hypothesis that incorporates sex differences in selection can contribute to sex chromosome differentiation, particularly when it occurs alongside other processes of sex chromosome degeneration that are currently recognized by evolutionary biologists. An appeal of this extended sheltering hypothesis is its simplicity and compatibility with several features of sex chromosome evolution, including recently discovered cases of X-linked gene losses that are not as easily explained by alternative scenarios for sex chromosome gene loss (see Table 1 for a comparison of predictions of the different models). Evaluating the broader importance of sheltering in sex chromosome evolution will, however, require renewed effort to infer the distribution of dominance for deleterious mutations and the sex-specific fitness costs of gene losses, which are critical parameters in these models. While these empirical aims will be challenging, modern genetic engineering approaches coupled with high throughput phenotyping place us in a good position to accomplish these goals.

**Table 1.**
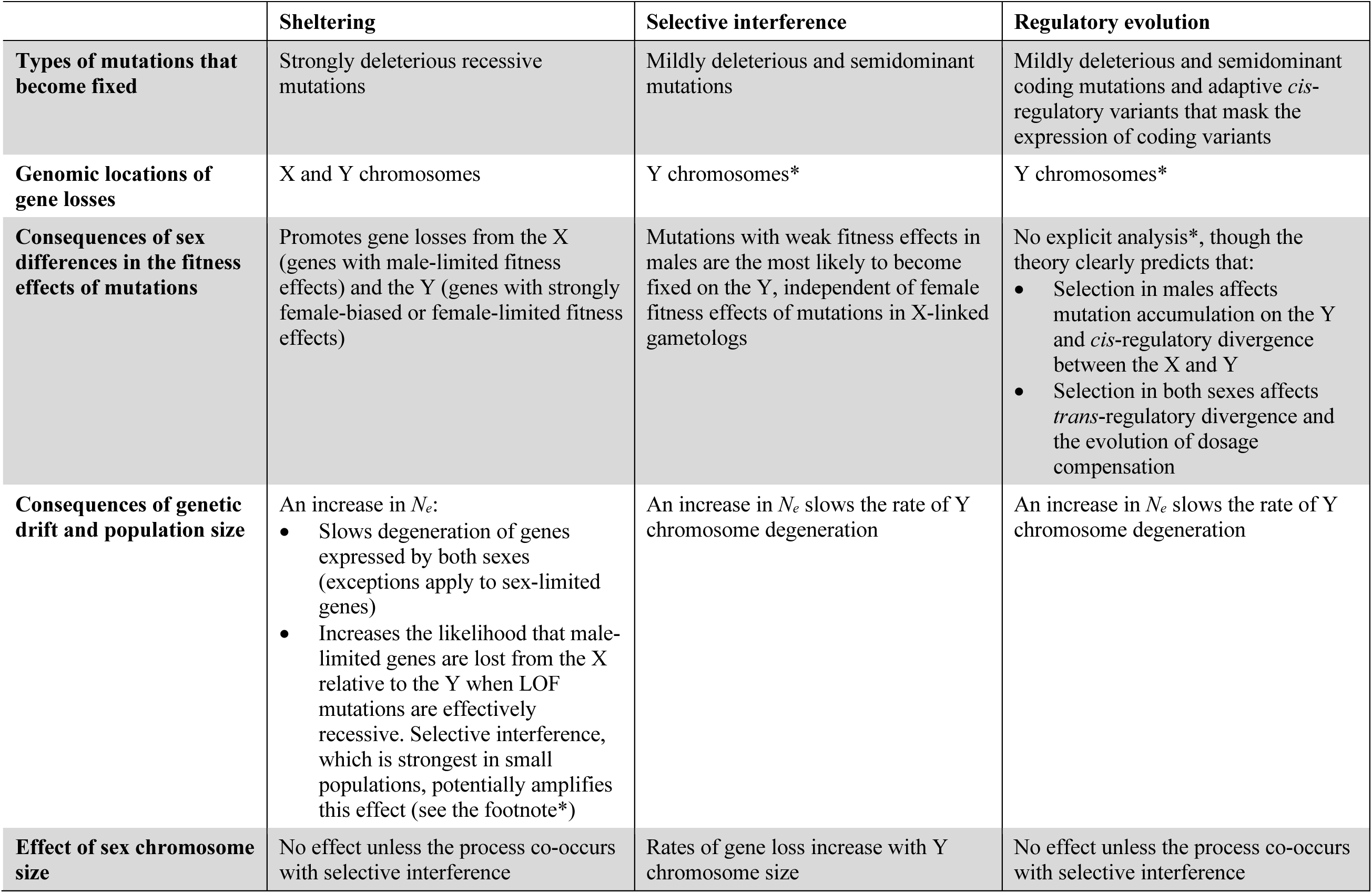
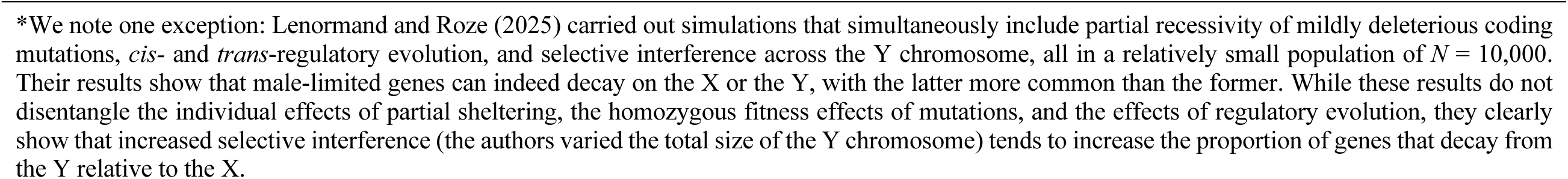
Summary of the predictions of different scenarios of sex chromosome gene loss.

|  | **Sheltering** | **Selective interference** | **Regulatory evolution** |
| --- | --- | --- | --- |
| **Types of mutations that become fixed** | Strongly deleterious recessive mutations | Mildly deleterious and semidominant mutations | Mildly deleterious and semidominant coding mutations and adaptive *cis*-regulatory variants that mask the expression of coding variants |
| **Genomic locations of gene losses** | X and Y chromosomes | Y chromosomes* | Y chromosomes* |
| **Consequences of sex differences in the fitness effects of mutations** | Promotes gene losses from the X (genes with male-limited fitness effects) and the Y (genes with strongly female-biased or female-limited fitness effects) | Mutations with weak fitness effects in males are the most likely to become fixed on the Y, independent of female fitness effects of mutations in X-linked gametologs | No explicit analysis*, though the theory clearly predicts that:<br>• Selection in males affects mutation accumulation on the Y and *cis*-regulatory divergence between the X and Y<br>• Selection in both sexes affects *trans*-regulatory divergence and the evolution of dosage compensation |
| **Consequences of genetic drift and population size** | An increase in $N_e$:<br>• Slows degeneration of genes expressed by both sexes (exceptions apply to sex-limited genes)<br>• Increases the likelihood that male-limited genes are lost from the X relative to the Y when LOF mutations are effectively recessive. Selective interference, which is strongest in small populations, potentially amplifies this effect (see the footnote*) | An increase in $N_e$ slows the rate of Y chromosome degeneration | An increase in $N_e$ slows the rate of Y chromosome degeneration |
| **Effect of sex chromosome size** | No effect unless the process co-occurs with selective interference | Rates of gene loss increase with Y chromosome size | No effect unless the process co-occurs with selective interference |

\*We note one exception: Lenormand and Roze (2025) carried out simulations that simultaneously include partial recessivity of mildly deleterious coding mutations, *cis*- and *trans*-regulatory evolution, and selective interference across the Y chromosome, all in a relatively small population of $N = 10,000$ . Their results show that male-limited genes can indeed decay on the X or the Y, with the latter more common than the former. While these results do not disentangle the individual effects of partial sheltering, the homozygous fitness effects of mutations, and the effects of regulatory evolution, they clearly show that increased selective interference (the authors varied the total size of the Y chromosome) tends to increase the proportion of genes that decay from the Y relative to the X.

## Acknowledgements

We thank Filip Ruzicka, Colin Olito, Akane Uesugi, Melissa Toups, and Daniel Jeffries for comments on an early version of the paper. We are particularly grateful to Deborah and Brian Charlesworth for commenting on two drafts of the manuscript. We thank Aneil Agrawal and Thomas Lenormand for email correspondence about the data on dominance and ways to interpret it. Technical support was provided by ISTA Scientific Computing Unit.

## Data Availability

Full details of the mathematical models on which the study is based are provided in the Supporting Information document associated with this article.

## Notes

### Competing Interest Statement

The authors have declared no competing interest.

