## Supporting Information for "A re-evaluation of Muller’s sheltering hypothesis for the evolution of sex chromosome gene content"

**Contents:**


- Appendix 1. Deterministic model of sheltering
- Appendix 2: Sheltering under mutation, selection and genetic drift
- Appendix 3: Hitchhiking and background selection models
- Appendix 4: Additional results for cases of incomplete recessivity
- Supplementary Figures
- Supplementary Methods
- Supplementary References

#### Appendix 1. Deterministic model of sheltering

We focus on genes present in functional and non-functional states on a pair of X and Y chromosomes that do not recombine with one another. Each gene has two major alleles: a functional allele that is favoured by natural selection and a deleterious loss-of-function (LOF) allele. Mutations in each gene are unidirectional, with functional alleles mutating to LOF alleles at rates of $\mu_f$ in female gametes and $\mu_m$ in male gametes. Gain of function mutations are assumed to be sufficiently rare that they can be ignored.

At a given gene, $A$ will be the functional allele and $A_0$ will be the LOF allele. Let $p_m$ and $p_f$ represent the frequencies (respectively) of the X-linked LOF allele in male and female gametes contributing to fertilization; $p_Y$ is the frequency of the same allele on Y-bearing male gametes contributing to fertilization. The corresponding functional allele frequencies are $q_m = 1 - p_m$, $q_f = 1 - p_f$, and $q_Y = 1 - p_Y$. Female and male gametes combine randomly to form zygotes of the next generation. Table S1 presents the zygotic frequencies and fitness per genotype. Selection and mutation complete a full generation cycle, at which point the frequency of the *A* allele in X-bearing male gametes will be:

$$1 - p_m' = \frac{(1-p_f)(1-p_Y s_m h)(1-\mu_m)}{p_f\big(1 - s_m h - p_Y s_m (1-h)\big) + (1-p_f)(1-p_Y s_m h)} \qquad [1a]$$

The frequency of the *A* allele in female gametes will be:

$$1 - p_f' = \frac{\frac{1}{2}\Big(p_f(1-p_m) + p_m(1-p_f)\Big)(1-s_f h) + (1-p_f)(1-p_m)}{1 - s_f p_f p_m - \Big(p_f(1-p_m) + p_m(1-p_f)\Big) s_f h}(1-\mu_f) \qquad [1b]$$

And the frequency of *A* on *Y*-bearing gametes will be:

$$1 - p_Y' = \frac{(1-p_Y)(1-p_f s_m h)(1-\mu_m)}{p_Y\Big(1 - s_m h - p_f s_m (1-h)\Big) + (1-p_Y)(1-p_f s_m h)} \qquad [1c]$$

Eqs. [1a-c] are exact and serve as the basis for all of our deterministic predictions.

**Table S1.** Frequency and fitness for each genotype and sex[1]

| | Genotype | | |
|---|---|---|---|
| | $A_0A_0$ | $A_0A$ | $AA$ |
| **Frequency in female zygotes** | $p_fp_m$ | $p_fq_m + p_mq_f$ | $q_fq_m$ |
| **Frequency in male zygotes** | $p_fp_Y$ | $p_fq_Y + p_Yq_f$ | $q_fq_Y$ |
| **Fitness in females** | $1 - s_f$ | $1 - s_fh$ | 1 |
| **Fitness in males** | $1 - s_m$ | $1 - s_mh$ | 1 |

[1] Note that males have two possible orientations of the heterozygous genotype: (*i*) heterozygous with an X-linked LOF allele, which occurs with frequency $p_fq_Y$, or (*ii*) heterozygous with a Y-linked LOF allele, which occurs with frequency $p_Yq_f$.

**Evolutionary dynamics and equilibria for recessive LOF alleles**

For cases where deleterious mutations are recessive ($h$ = 0), and mutation rates are positive, the recursions simplify to:

$$p_m' = 1 - \frac{(1 - p_f)(1 - \mu_m)}{1 - s_m p_Y p_f}$$

$$p_f' = 1 - \frac{1 - \frac{1}{2}(p_f + p_m)}{1 - s_f p_f p_m}(1 - \mu_f)$$

$$p_Y' = 1 - \frac{(1 - p_Y)(1 - \mu_m)}{1 - s_m p_Y p_f}$$

In this case, there are four possible equilibria:

1. Fixation on the X and Y ($\hat{p}_Y = \hat{p}_f = \hat{p}_m = 1$)
2. Fixation on the Y and polymorphism on the X ($\hat{p}_Y = 1; \hat{p}_f < 1; \hat{p}_m < 1$)
3. Fixation on the X and polymorphism on the Y ($\hat{p}_Y < 1; \hat{p}_f = \hat{p}_m = 1$)
4. Polymorphism on both the X and Y ($\hat{p}_Y < 1; \hat{p}_f < 1; \hat{p}_m < 1$)

The fourth equilibrium state, when valid, is:

$$\hat{p}_X = \hat{p}_m = \hat{p}_f = \sqrt{\frac{\mu_f}{s_f}} \qquad [2a]$$

$$\hat{p}_Y = \frac{\mu_m}{s_m \hat{p}_X} = \frac{\mu_m}{s_m}\sqrt{\frac{s_f}{\mu_f}} \qquad [2b]$$

The X-linked equilibrium will be higher than the Y-linked equilibrium when:

$$\hat{p}_X > \hat{p}_Y \Leftrightarrow \frac{s_m}{s_f} > \frac{\mu_m}{\mu_f}$$

The X-linked equilibrium will be higher than the Y-linked equilibrium when:

$$\hat{p}_X < \hat{p}_Y \Leftrightarrow \frac{s_m}{s_f} < \frac{\mu_m}{\mu_f}$$

The equilibrium with polymorphism on the X and Y will be valid under the condition: $\mu_m/s_m < \sqrt{\mu_f/s_f} < 1$, which defines the minimum requirements of selection each sex to prevent degeneration on either chromosome. With equal mutation rates and selection in each sex ($s_f = s_m$; $\mu_f = \mu_m$), this equilibrium further simplifies to:

$$\hat{p}_f = \hat{p}_m = \hat{p}_Y = \sqrt{\frac{\mu}{s}}$$

which matches Fisher's (1935) result (*22*), in which harmful mutations (lethal alleles in his model: $s$ = 1) evolve to equal frequencies on the X and Y chromosome.

**Deterministic fixation of LOF alleles on the X**

The X-linked LOF allele will be driven to fixation when $s_f \leq \mu_f$, which should apply to male-limited genes (genes with no effect on female fitness: $s_f = 0$). This result is implied by the equilibrium in eq. [2a], and it can be formally evaluated by way of a linear stability analysis for the equilibrium in which the LOF is fixed on the X and the Y evolves to a polymorphic equilibrium at mutation-selection balance. This equilibrium corresponds to: $\hat{p}_f = \hat{p}_m = 1$ and $\hat{p}_Y = \mu_m(1 - hs_m)/s_m(1 - h)$. The characteristic polynomial for this equilibrium (by way of the Jacobian matrix for the set of exact recursion equations; see chapter 8 of Otto and Day 2007) is:

$$\det\begin{pmatrix} -\lambda & 1 & 0 \\ \frac{(1-\mu_f)}{2(1-s_f)} & \frac{(1-\mu_f)}{2(1-s_f)}-\lambda & 0 \\ 0 & 1-\frac{\mu_m}{s_m} & \frac{1-s_m}{1-\mu_m}-\lambda \end{pmatrix}$$

$$= \left(\lambda^2 - \lambda\frac{(1-\mu_f)}{2(1-s_f)} - \frac{(1-\mu_f)}{2(1-s_f)}\right)\left(\frac{1-s_m}{1-\mu_m}-\lambda\right) = 0$$

which yields the eigenvalues:

$$\lambda = \frac{1-s_m}{1-\mu_m}$$

$$\lambda = \frac{\frac{(1-\mu_f)}{2(1-s_f)} - \sqrt{\left(\frac{(1-\mu_f)}{2(1-s_f)}\right)^2 + 4\frac{(1-\mu_f)}{2(1-s_f)}}}{2}$$

$$\lambda = \frac{\frac{(1-\mu_f)}{2(1-s_f)} + \sqrt{\left(\frac{(1-\mu_f)}{2(1-s_f)}\right)^2 + 4\frac{(1-\mu_f)}{2(1-s_f)}}}{2}$$

Provided $\mu_m < s_m$ (which must be true for $\hat{p}_Y < 1$ given $h = 0$), then the first eigenvalue is always less than one. Because $\frac{(1-\mu_f)}{2(1-s_f)} > 0$, the third eigenvalue is the one we need to focus on. The equilibrium will be stable when that eigenvalue is less than one, *i.e.*:

$$1 > \frac{\frac{(1-\mu_f)}{2(1-s_f)} + \sqrt{\left(\frac{(1-\mu_f)}{2(1-s_f)}\right)^2 + 4\frac{(1-\mu_f)}{2(1-s_f)}}}{2}$$

which simplifies to $s_f < \mu_f$. The same eigenvalue is greater than one—and LOF allele fixation will therefore be unstable—when $s_f > \mu_f$.

**Deterministic fixation of LOF alleles on the Y**

The Y-linked LOF allele will be driven to fixation when $s_m \le \alpha\sqrt{\mu_f s_f}$, where $\alpha = \mu_m/\mu_f$. The condition for stable fixation is implied by eq. [2b], and a stability analysis of the equilibrium with $p_Y = 1$ confirms the prediction. Specifically, when the LOF allele is near fixation on the Y and the *A* allele is near fixation on the X (the latter is expected when selection in females is strong relative to the mutation rate), then the X-linked dynamics are approximately:

$$p'_X = p_X + \frac{-2s_f p_X^2 - s_m p_Y p_X + 2\mu_f + \mu_m}{3}$$

which yields the following X-linked equilibrium when the LOF allele is fixed on the Y:

$$\hat{p}_X = \frac{\sqrt{s_m^2 + 8s_f(2\mu_f + \mu_m)} - s_m}{4s_f}$$

Deterministic simulations using the exact recursions show that the equilibrium is accurate in cases where selection in females is strong relative to the mutation rate. The following figure compares the approximate equilibrium (grey) against exact forward simulations to equilibrium using the full and exact recursion equations (red) (results use $\mu_f = \mu_m = 10^{-4}$ and $s_f = 0.1$).

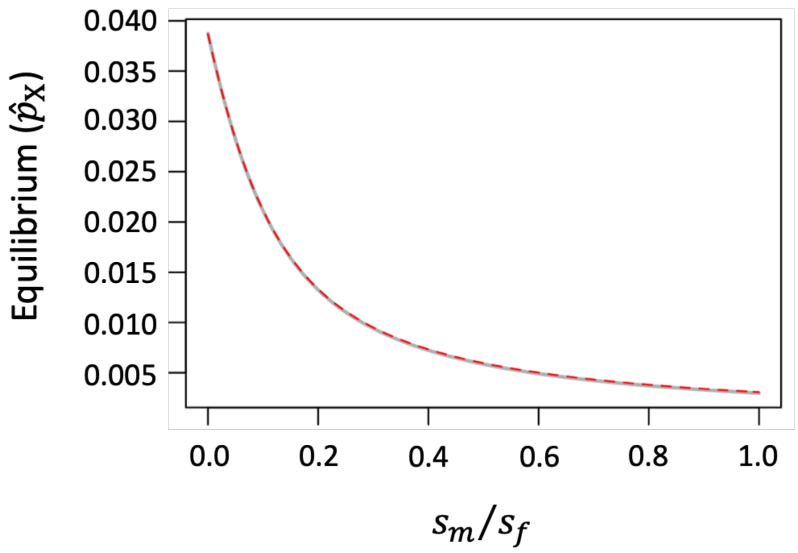


Using the approximate recursions, we obtain the following Jacobian matrix for the equilibrium with Y-linked LOF alleles fixed:

$$J = \begin{pmatrix} 1 - \frac{4s_f\hat{p}_X + s_m}{3} & -\frac{s_m\hat{p}_X}{3} \\ 0 & \frac{1-\mu_m}{1-s_m\hat{p}_X} \end{pmatrix}$$

which has the leading eigenvalue:

$$\lambda_L = \frac{1-\mu_m}{1-s_m\hat{p}_X}$$

The equilibrium will be stable if $\lambda_L < 1$, which requires:

$$s_m < \frac{\mu_m}{\hat{p}_X}$$

Substituting the approximation for $\hat{p}_X$, we have:

$$s_m < \alpha\sqrt{\mu_f s_f}$$

where $\alpha = \mu_m/\mu_f$, which confirms the same result stated above.

It is clear from this last result that conditions for fixation of LOF alleles are more permissive on the Y chromosome than they are on the X. Recall that on the X chromosome, fixation was only likely for genes that have negligible importance in females. In contrast, deterministic gene degeneration can occur on the Y chromosome in cases where the gene has a meaningful fitness effect in males. For example, for genes that are essential for females ($s_f$ = 1) and that have relatively large LOF mutation rates ($\mu_f = \mu_m = 10^{-4}$), LOF mutations on the Y will deterministically spread to fixation when $s_m < 0.01$ (*i.e.*, mutations altering fitness by up to 1%). Male-biased mutation rates will further expand the criteria for deterministic degeneration. This effect arises because strong selection in females keeps LOF alleles on the X at low frequencies, which generates strong sheltering of Y-linked genes in males.

**General dynamics of sex-limited genes**

For genes with female-limited functions ($s_f > 0 = s_m$), the Y-linked recursion simplifies to:

$$1 - p'_Y = (1 - p_Y)(1 - \mu_m)$$

which is independent of selection in females and allows for a general solution of the allele frequency dynamics on the Y. The expected frequency of Y-linked LOF mutations after $t$ generation of evolution will be:

$$p_{Y,t} = 1 - \left(1 - p_{Y,0}\right)(1 - \mu_m)^t \approx 1 - \left(1 - p_{Y,0}\right)e^{-\mu_m t}$$

where $p_{Y,0}$ is the initial frequency; the final approximation applies because $\mu_m$ is small. If we assume that the LOF allele was initially absent from the population ($p_{Y,0} = 0$), the result further simplifies to $p_{Y,t} \approx 1 - e^{-\mu_m t}$, as presented in the main text. For the mirror image case involving LOF mutations in male-limited genes, we can approximate the evolutionary dynamics on the X and Y chromosome as:

$$p_X' \approx p_X + \frac{-s_m p_Y p_X(1 - p_X) + \left(\mu_m + 2\mu_f\right)(1 - p_X)}{3}$$

$$p_Y' \approx p_Y - s_m p_X p_Y(1 - p_Y) + \mu_m(1 - p_Y)$$

These dynamics suggest that the Y chromosome should respond more quickly to selection (by a factor of three) than the X. We may therefore use a separation of timescales approximation, in which we assume that the Y-linked allele frequencies will quickly approach the following a quasi-equilibrium state given the frequency of the X-linked mutation:

$$\tilde{p}_Y \approx \frac{\mu_m}{s_m p_X}$$

Plugging this quasi-equilibrium into the expression for change $p_X$ gives us:

$$1 - p_X' \approx (1 - p_X)\left(1 - \frac{2\mu_f}{3}\right)$$

which yields the general solution:

$$p_{X,t} \approx 1 - \left(1 - p_{X,0}\right)\left(1 - \frac{2\mu_f}{3}\right)^t \approx 1 - \left(1 - p_{X,0}\right)\exp\left(-\frac{2}{3}\mu_f t\right)$$

where $p_{X,0}$ is the initial X-linked frequency of the LOF allele, and $p_{X,t}$ is its frequency after $t$ generations. With the X-linked LOF initially absent from the population, the last result further simplifies to $p_{X,t} \approx 1 - \exp\left(-\frac{2}{3}\mu_f t\right)$, as presented in the main text.

**Deterministic changes in the genetic load**

The equilibrium genetic load contributed by a locus mutating to recessive LOF alleles is define as follows. For an autosomal locus—which includes the ancestral state preceding the origin of a new sex chromosome system or a translocation event that generates neo-X and neo-Y chromosomes—the female and male genetic loads will be:

$$L_f = \hat{p}_m \hat{p}_f s_f$$

and

$$L_m = \hat{p}_m \hat{p}_f s_m$$

where $\hat{p}_f$ and $\hat{p}_m$ are the equilibrium frequencies of LOF alleles in female and male gametes contributing to fertilization in each generation.

For an autosomal locus that mutates to recessive LOF alleles with homozygous fitness effects of $s_m$ in males and $s_f$ in females, the evolutionary dynamics under mutation and selection will be:

$$1 - p'_m = \left(1 - \frac{p_m p_f (1 - s_m) + \frac{1}{2}(p_f + p_m - 2p_f p_m)}{1 - p_m p_f s_m}\right)(1 - \mu_m)$$

$$1 - p'_f = \left(1 - \frac{p_m p_f (1 - s_f) + \frac{1}{2}(p_f + p_m - 2p_f p_m)}{1 - p_m p_f s_f}\right)(1 - \mu_f)$$

Note that these frequency dynamics should also be roughly equivalent to those of pseudo-autosomal regions (PAR) provided the locus in question is not closely linked to the male-limited region of the Y. At equilibrium for this model, we have the identity:

$$2 = \frac{1-\mu_f}{1-\hat{p}_m\hat{p}_f s_f} + \frac{1-\mu_m}{1-\hat{p}_m\hat{p}_f s_m}$$

For female-limited loci, the equilibrium load (in females) simplifies to:

$$L_f = \hat{p}_m\hat{p}_f s_f = \frac{\mu_f + \mu_m}{1+\mu_m} = \mu_f + \frac{\mu_m(1-\mu_f)}{1+\mu_m}$$

And for male-limited loci, the equilibrium load (in males) becomes:

$$L_m = \hat{p}_m\hat{p}_f s_m = \frac{\mu_m + \mu_f}{1+\mu_f} = \mu_m + \frac{\mu_f(1-\mu_m)}{1+\mu_f}$$

What happens to the load at equilibrium for genes on sex chromosomes, particularly genes that have become degenerate on either the X or Y? We focus on the bookend cases of our deterministic model, which include male-limited gene degeneration is expected on the X chromosome, and female-limited gene degeneration is expected on the Y.

For a female-limited gene, the Y-linked copy is expected to degenerate while the X-linked copy will evolve to the equilibrium at mutation-selection balance. In this case, the equilibrium for the X is approximately:

$$\hat{p}_X = \sqrt{\frac{2\mu_f + \mu_m}{2s_f}}$$

and the equilibrium load for females becomes:

$$L_f = s_f\hat{p}_X^2 \approx \mu_f + \frac{\mu_m}{2}$$

which is less than the ancestral autosomal load as long as $1 > \mu_m + 2\mu_f$. This condition will always be true given biologically realistic mutation rates.

For a male-limited gene, the X-linked copy is expected to degenerate while the Y-linked copy will evolve to the equilibrium at mutation-selection balance. In this case, the equilibrium for the Y is:

$$\hat{p}_Y = \frac{\mu_m}{s_m}$$

and the male load will be:

$$L_m = s_m \hat{p}_Y = \mu_m$$

which is again an improvement over the ancestral autosomal state provided $\mu_f > 0$.

Finally, consider the intermediate case of a gene in which LOF mutations equally affect the sexes. Although, sheltering in the absence of drift is not expect to cause degeneration in this case, the transition from an ancestral autosomal state to a sex-linked state does have an effect on the equilibrium genetic load. For an autosome with $s = s_f = s_m$, the equilibrium load at mutation-selection balance is:

$$L_f = L_m = \hat{p}_m \hat{p}_f s = \frac{\mu_f + \mu_m}{2}$$

In cases where both the X and Y chromosome equilibria are polymorphic, which includes the specific case of equal selection in each sex (see above), the female and male genetic loads become:

$$L_f = s_f \hat{p}_f \hat{p}_m = \mu_f$$

$$L_m = s_m \hat{p}_f \hat{p}_Y = \mu_m$$

The autosomal and sex-linked loads will all be the same in the special case where mutation rates show no sex bias. If, as in many species, the mutation rate is male-biased ($\mu_m > \mu_f$), evolutionary transition from an autosomal to a sex-linked state will result in a reduction of the female load and elevation of the male load.

#### Appendix 2: Sheltering under mutation, selection and genetic drift

We incorporate drift into our models using a combination of exact computer simulations of X- and Y-linked evolutionary dynamics in a Wright-Fisher population where multinomial sampling of genotypes mimics the process of genetic drift (see pp. 229-230 of Charlesworth and Charlesworth 2010). Specifically, we assume in each generation that there are a fixed number of adult females and males ($N_f$ and $N_m$, respectively, though we focus on the simplest case where $N_f = N_m$). The deterministic model presented in Appendix 1 yields predictions for the expected frequencies of each female and male genotype in the post-selection pool of individuals contributing to reproduction. We then carry out independent multinomial sampling in each sex (using a pseudo-random number generator) to determine the actual genotype frequencies in the pool of reproducing adults. Mutations occur after the multinomial sampling step and cause some deviation between LOF allele frequencies of adults and the allele frequencies in gametes contributing to fertilization in the next generation. Given an equal sex ratio, the effective population size for autosomes will be $2N_e$, where $N_e = N_f + N_m$; Effective sizes for the X and Y will be $1.5N_e$ and $0.5N_e$, respectively.

***Y-linked fixation probabilities.*** Each of our simulations of Y-linked fixation probabilities (as in Fig. 2) begin with a single initial copy of a Y-linked LOF allele. The initial frequency of the X-linked LOF allele corresponds to the mean of the stationary distribution predicted for the X chromosome in an ancestral population in which all Y-linked copies were functional (the initial condition is derived immediately below). The system was then allowed to evolve under recurrent mutation on the X (but not the Y), and selection and drift on both the X and Y, until the Y-linked variant is either lost from the population or fixed. The proportion of fixations among the set of simulation runs was used to calculate the fixation probability of a Y-linked mutation.

The starting conditions for these fixation probability simulations require some further explanation. While the initial conditions allow us to make clear conclusions about the evolutionary fates of unique Y-linked variants entering the population, the predictions they yield should be conservative (they should underestimate actual fixation probabilities on the Y) and therefore interpreted as such. The initial state is of a population where X-linked LOF alleles will, if anything, be artificially high because we leave no opportunity for selection on males to influence the initial X-linked variability. Thus, once we introduce a new Y-linked variant, the sheltering effect it experiences will be dampened relative to a population in which both X and Y segregate at mutation-selection balance. This is why fixation probabilities can be less than that of a neutral mutation in scenarios in which our deterministic model predicts their fixation.

***Simulations of X versus Y chromosome degeneration.*** Our models exploring the rates and relative probabilities of gene degeneration on the X and Y (as in Fig. 3) are based on the full model that includes selection, drift, and recurrent mutation at both X-linked and Y-linked loci. For these simulations, our initial population is fixed at both chromosomes for the functional allele. From this initial state, we carried out stochastic simulations until a LOF allele is fixed on either the X or the Y and we recorded the outcome and the dynamical trajectory of the allele that reaches fixation.

All simulations were carried out in R (R Core Team. 2021).

**Fixation probabilities of new Y-linked mutations: analytical approximations**

Our analytical approach follows that of Nei (*23*), who modelled the fixation probabilities for Y-linked mutations entering a population that was polymorphic for the X. We will assume here that purifying selection on the X is strong relative to the mutation rate, which ensures that X-linked LOF mutations will be rare. The requirement is easily met as long as selection in females is strong relative to the mutation rate, and the population scaled selection coefficient ($N_e s_f$) is large. We will also ignore X-linked allele frequency differences between sexes, which will be negligible under the preceding assumption that X-linked purifying selection is strong. We focus on recessive LOF alleles.

In ancestral population with no Y-linked variation, males cannot be homozygous, whereas females will sometimes be homozygous for X-linked alleles. Thus, all ancestral purifying selection that governs the initial diversity on the X is due to selection on females. As in Nei (*23*), we assume that the initial X-linked diversity is at equilibrium between mutation, purifying selection, and genetic drift.

In the ancestral population, the expected change in X-linked LOF alleles, per generation, is:

$$M_X \approx -\frac{2}{3}s_f p_X^2(1-p_X) + \frac{2\mu_f + \mu_m}{3}(1-p_X)$$

which is a function of selection in females and mutation in both sexes. Assuming that X-linked genes have three-quarters effective population size of autosomes ($2N_e$ for autosomes; $1.5N_e$ for the X), then the variance in X-linked allele frequency change, per generation, is:

$$V_X = \frac{p_X(1-p_X)}{1.5N_e}$$

Given these approximations, and applying the standard diffusion approximation for mutation-selection-drift balance (*e.g.*, Wright 1945; Crow and Kimura 1970), the stationary distribution for X-linked LOF mutations will be:

$$f(p_X) = \frac{C}{V}\exp\left(2\int\frac{M_X}{V_X}dp\right) = Cp_X^{N_e(2\mu_f+\mu_m)-1}(1-p_X)^{-1}e^{-N_e s_f p_X^2}$$

where $C$ is a constant that ensures that the distribution integrates to one. Following Nei (1968), and assuming that $N_e s_f$ is large so that the terms $(1-p_X)^{-1}$ can be ignored, then the mean of the stationary distribution can be approximated as:

$$\bar{p}_X = \frac{\Gamma\left(\frac{1}{2}N_e(2\mu_f + \mu_m) + \frac{1}{2}\right)}{\sqrt{N_e s_f}\,\Gamma\left(\frac{1}{2}N_e(2\mu_f + \mu_m)\right)}$$

where $\Gamma(x)$ is the gamma function. In the special case where $\mu = \mu_f = \mu_m$, the last result simplifies to

$$\bar{p}_X = \frac{\Gamma\left(1.5N_e\mu + \frac{1}{2}\right)}{\sqrt{N_e s_f}\,\Gamma(1.5N_e\mu)}$$

which mirrors eq. (4) of Nei (*23*), but with one exception. In our model we use the term $\sqrt{N_e s_f}$ rather than $\sqrt{1.5N_e s}$ (as in (*23*)) because purifying selection is female-limited owing to the absence of segregating LOF alleles on the Y (Nei's model implies that purifying selection can occur in males as well, but this is not possible under the assumption that the ancestral Y is monomorphic for functional alleles). The following figure plots our expression for $\bar{p}_X$ as a function of $N_e\mu$ (solid curve) and compares it to the deterministic mutation-selection equilibrium, $\hat{p}_X = \sqrt{(2\mu_f + \mu_m)/2s_f}$ (broken line). The latter is a reasonable approximation of the former when $N_e\mu \gg 1$. Smaller population-scaled mutation rates result in mean LOF allele frequencies that are lower than deterministic.

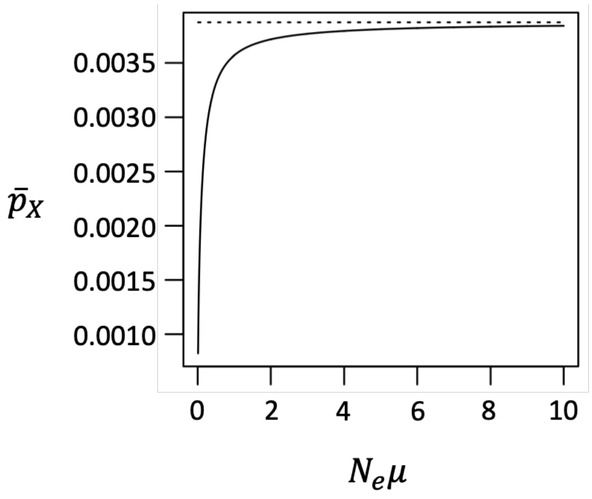

Given an X-linked LOF allele frequency of $\bar{p}_X$, the expected allele frequency change on the Y chromosome (ignoring additional mutations) is:

$$M_Y \approx -s_m \bar{p}_X p_Y (1 - p_Y)$$

The variance in Y-linked allele frequency change per generation is $V_Y = 2p_Y(1 - p_Y)/N_e$. Following Kimura (1962) and Nei (1970), the fixation probability for a Y-linked mutation with initial frequency $p_0$ will be:

$$U_Y(p_0) = \frac{\int_0^{p_0} G(x)dx}{\int_0^1 G(x)dx} = \frac{e^{N_e s_m \bar{p}_X p_0} - 1}{e^{N_e s_m \bar{p}_X} - 1}$$

where:

$$G(x) = \exp\left(-2\int_0^x \frac{M_Y}{V_Y} dp_Y\right) = \exp(N_e s_m \bar{p}_X x)$$

With $0.5 N_e \mu_m$ new Y-linked mutations per generation, each with initial frequency $p_0 = 2/N_e$, then the rate of fixation for Y-linked LOF alleles should be:

$$R_Y = \frac{1}{2} N_e \mu_m U_Y(p_0) = \frac{1}{2} N_e \mu_m \frac{e^{2 s_m \bar{p}_X} - 1}{e^{N_e s_m \bar{p}_X} - 1}$$

The following figure plots fixation probabilities of Y-linked LOF mutations in genes that are essential for females ($s_f = 1$). Two different population sizes and three LOF mutation rates are considered ($\mu = (2\mu_f + \mu_m)/3$). The results show that sheltering promotes Y-linked degeneration in cases where the gene is much less important in males than females ($s_m << s_f$), effective population size is small, and LOF mutation rates are low (*e.g.,* small genes). The latter effect is attributable to the fact that $\bar{p}_X$ declines relative to deterministic predictions with declining $N_e \mu$ (see the preceding figure), which enhances the masking effect the X chromosome has on Y-linked LOF alleles.

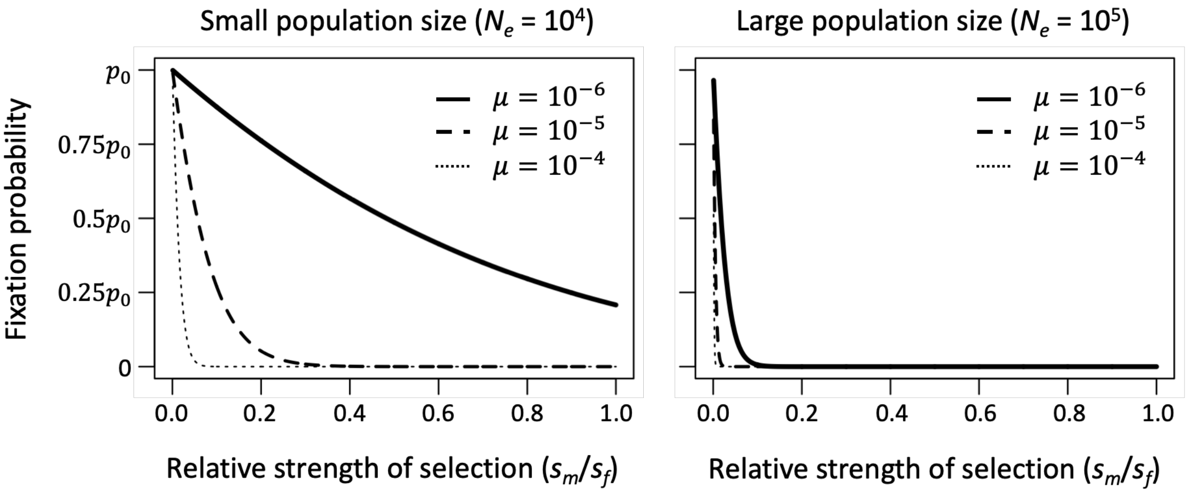


Our diffusion approximations, like those of Nei (*23*), predict fixation probabilities at Y-linked genes while holding the X-linked LOF allele frequency constant and equal to the mean predicted from the stationary distribution for the X. In other words, the following approximations neglect evolution on the X chromosome following the introduction of new genetic variants on the Y. This is obviously an unrealistic assumption, but a necessity for any analytical progress with the model. We therefore present the approximations in combination with full stochastic forward simulations that relax this critical assumption by allowing the X and Y to dynamically co-evolve with one another. The simulations suggest that the analytical approximation is most accurate when $N_e\mu$ is large, and it otherwise underestimates the true fixation probability (see Fig. 2 in the main text), presumably because drift often causes deleterious allele on the X to drop sharply enough that the that sheltering of Y-linked effects become amplified and increase the likelihood that the Y-linked allele then becomes fixed.

***Evolutionary fates of male-limited genes in small populations.*** In large populations, male-limited genes are expected to degenerate on the X chromosome while being retained on the Y. However, as the population declines and the population-scaled mutation rate becomes small, sheltering effects become stronger on both types of chromosomes because LOF alleles become so rare. New X-linked LOF variants become strongly sheltered because the Y tends to be fixed for the functional copy. Likewise, Y-linked variants become strongly sheltered because the X also tends to be fixed for the functional copy. In the limit of mutation-limited evolution, where $N_e\mu \rightarrow 0$, the X and Y chromosomes essentially compete to contribute the first LOF allele fixation. Whichever chromosome degenerates first for the male-limited gene, will cause effectively strong selection on the other chromosome to maintain the function of the same gene. And because the population is largely homomorphic prior to the fixation event, the evolutionary dynamics of each male-limited LOF allele that enters the population will be dominated by drift. Such alleles will tend to be either lost or fixed before the next LOF mutation enters the population.

In this mutation-limited environment, the probability that a new X-linked variant arises and is ultimately fixed will be $R_X = (2\mu_f + \mu_m)/3$ per generation. The corresponding probability on the Y will be $R_Y = \mu_m$. The probability that the first fixation event is on the X will be:

$$\frac{R_X}{R_X + R_Y} = \frac{2 + \mu_m/\mu_f}{2(1 + 2\,\mu_m/\mu_f)}$$

which corresponds to the limit presented in the main text (e.g., see Fig. 3a).

#### Appendix 3: Hitchhiking and background selection models

**Fixation of LOF alleles via hitchhiking: results for a single Y-linked gene**

***Exact deterministic dynamics of the hitchhiking model.*** Assume that are two alleles at the X-linked locus (the functional allele *A* and a LOF allele $a$) and *effectively* three alleles on the Y (owing to complete linkage): the functional allele, a LOF allele in an otherwise ancestral Y chromosome background, and a LOF allele on a background with a beneficial mutation. We assume that the beneficial allele increases male fitness by a factor of $1 + s_b$, and fitness effects across loci are multiplicative. The LOF allele frequencies on the X are represented by $p_m$ and $p_f$ (as before), the frequency of the LOF allele on the ancestral background is $p_Y$, and the frequency of the LOF/beneficial combination is $p_b$. The evolutionary dynamics are described by the following recursions.

The frequency of the *A* allele in X-bearing male gametes will be:

$$1 - p'_m = \frac{(1 - p_f)\big((1 - p_Y - p_b) + p_Y(1 - s_m h) + p_b(1 - s_m h)(1 + s_b)\big)(1 - \mu_m)}{\bar{w}_m}$$

where:

$$\bar{w}_m = (1 - p_f)(1 - p_Y - p_b) + p_Y(1 - p_f)(1 - s_m h) + p_b(1 - p_f)(1 - s_m h)(1 + s_b) + p_f(1 - p_Y - p_b)(1 - s_m h) + p_Y p_f(1 - s_m) + p_b p_f(1 - s_m)(1 + s_b)$$

The frequency of the *A* allele in female gametes will be:

$$1 - p'_f = \frac{\frac{1}{2}\Big(p_f(1 - p_m) + p_m(1 - p_f)\Big)(1 - s_f h) + (1 - p_f)(1 - p_m)}{1 - s_f p_f p_m - \Big(p_f(1 - p_m) + p_m(1 - p_f)\Big) s_f h}(1 - \mu_f)$$

The frequency of *A* on *Y*-bearing gametes will be:

$$1 - p'_Y - p'_b = \frac{(1 - p_Y)(1 - p_f s_m h)(1 - \mu_m)}{\bar{w}_m}$$

The frequency of the LOF allele on Y-bearing gametes <u>*without*</u> the beneficial mutation is:

$$p'_Y = \frac{p_Y\Big(1 - s_m h - p_f s_m(1 - h)\Big)}{\bar{w}_m} + \frac{\mu_m(1 - p_Y)(1 - p_f s_m h)}{\bar{w}_m}$$

The frequency of the LOF allele on Y-bearing gametes *with* the beneficial mutation is:

$$p_b' = \frac{p_b\left(1 - s_m h - p_f s_m(1-h)\right)(1+s_b)}{\overline{w}_m}$$

When LOF alleles are completely recessive ($h$ = 0), the system simplifies to:

$$p_m' = 1 - \frac{(1-p_f)(1+p_b s_b)(1-\mu_m)}{\overline{w}_m}$$

$$p_f' = 1 - \frac{\frac{1}{2}\left(p_f(1-p_m) + p_m(1-p_f)\right) + (1-p_f)(1-p_m)}{1 - s_f p_f p_m}(1-\mu_f)$$

$$p_Y' = \frac{p_Y(1-p_f s_m)}{\overline{w}_m} + \frac{\mu_m(1-p_Y)}{\overline{w}_m}$$

$$p_b' = \frac{p_b(1-p_f s_m)(1+s_b)}{\overline{w}_m}$$

$$\overline{w}_m = 1 - p_Y p_f s_m + p_b\left(s_b - p_f s_m(1+s_b)\right)$$

***Approximations for the fixation probability of a LOF allele.*** Whether or not a beneficial mutation becomes established on the Y is affected by the marginal fitness of Y chromosomes that carry the beneficial variant. Suppose a beneficial mutation arises on a Y chromosome that carries a LOF allele with homozygous fitness effects $s_f$ and $s_m$ in females and males, respectively. The marginal fitness of a Y chromosome with the pair of alleles is:

$$w_b = (1+s_b)(1-p_X) + (1+s_b)(1-s_m)p_X = (1+s_b)(1-s_m p_X)$$

The marginal fitness of a Y chromosome that carries neither allele is unity. Assume that the LOF mutation is initially at the deterministic equilibrium ($\hat{p}_X$ and $\hat{p}_Y$ on the X and Y, respectively). Provided there is a net beneficial effect of a Y haplotype carrying a new beneficial mutation ($w_b$ > 1), then the probability of establishment for a double-mutant haplotype will be:

$$\sim 2(w_b - 1) = 2(1+s_b)(1-s_m\hat{p}_X) - 2 \approx 2(s_b - s_m\hat{p}_X)$$

The approximation is valid when $s_b > s_m\hat{p}_X$ and the establishment probability is zero otherwise.

Supposing that $s_b$ follows an exponential distribution with mean $\bar{s}_b$, then the probability that a new beneficial mutation both arises in association with the LOF allele and establishes is:

$$\Pr(\text{LOF fixes}) \approx \hat{p}_Y \int_{s_m\hat{p}_X}^{\infty} 2(s_b - s_m\hat{p}_X)\frac{1}{\bar{s}_b}e^{-s_b/\bar{s}_b}ds_b = 2\bar{s}_b\hat{p}_Y \exp\left(-\frac{s_m\hat{p}_X}{\bar{s}_b}\right)$$

where $\bar{s}_b^{-1}e^{-s_b/\bar{s}_b}$ is the probability density function for beneficial mutation effects.

***Simulations.*** Assume that the beneficial mutation is initially absent and both the X and Y are at polymorphic equilibrium for the LOF allele. Assuming that LOF mutations are recessive, the equilibria are: $\hat{p}_X = \hat{p}_m = \hat{p}_f = \sqrt{\frac{\mu_f}{s_f}}$ and $\hat{p}_Y = \min\left(1, \frac{\mu_m}{s_m\hat{p}_X}\right)$. To calculate the probability that a single beneficial mutation arising on the Y results in a hitchhiking event that fixes the LOF allele, we first calculated the probability that the mutation arises in association with the LOF allele by sampling from a Bernoulli distribution where $\hat{p}_Y$ is probability of success. In cases the mutation was associated with the LOF allele, we carried out simulations with selection, mutation and drift until the beneficial mutation was fixed or lost from the population. For each simulation run, we sampled a selection coefficient for the beneficial mutation ($s_b$) from an exponential distribution with mean $\bar{s}_b$. Simulations used the initial allele frequencies: $p_f = p_m = \hat{p}_X$, $p_Y = \hat{p}_Y - N_m^{-1}$ and $p_b = N_m^{-1}$, where $N_m$ is the number of males in the population (we assume an equal sex ratio so that $N_m = N_f = N/2$, where $N$ is the effective population size).

Each generation of the simulation used exact deterministic selection equations to predict the expected frequencies of females and males after selection. The actual number of breeding adults with each genotype was based on multinomial sampling of genotypes whose sampling probabilities were based on deterministic predictions. The breeding population was comprised of $N_m$ and $N_f$ adult males and females. The fixation probability was estimated as the proportion of beneficial mutations that landed on a Y carrying a LOF allele and was then fixed.

**Fixation of LOF alleles via hitchhiking: multiple Y-linked genes**

With multiple functional genes on the Y, single hitchhiking events can potentially result in fixation of LOF alleles at multiple genes. For simplicity, suppose that there are $n$ Y-linked genes, each with the same mutation rate to LOF alleles and the same homozygous selection coefficients in each sex. The number of LOF alleles per Y chromosome will then be Poisson distributed with mean and variance $n\hat{p}_Y$. The fixation probability of a beneficial mutation with fitness effect $s_b$ that is associated with $k$ LOF alleles is $\sim 2(1+s_b)(1-s_m\hat{p}_X)^k - 2 \approx 2(s_b - s_m\hat{p}_X k)$, which is valid when $s_b > s_m\hat{p}_X k$ and the probability is zero otherwise. Assuming that beneficial effects are drawn from an exponential distribution, then the fixation probability of a random beneficial mutation that is initially associated with $k$ LOF alleles is:

$$\Pr(\text{fix}|k) \approx \int_{s_m\hat{p}_X k}^{\infty} 2(s_b - s_m\hat{p}_X k)\frac{1}{\bar{s}_b}e^{-s_b/\bar{s}_b}ds_b = 2\bar{s}_b \exp\left(-\frac{s_m\hat{p}_X k}{\bar{s}_b}\right)$$

The expected number of LOF mutations fixed for each new beneficial mutation entering the population is:

$$\mathrm{E}(k_{LOF}) = \sum_{k=0}^{n} k \Pr(\text{fix}|k)\frac{e^{-n\hat{p}_Y}(n\hat{p}_Y)^k}{k!} \approx 2\bar{s}_b \sum_{k=0}^{n} k \exp\left(-\frac{s_m\hat{p}_X k}{\bar{s}_b}\right)\frac{e^{-n\hat{p}_Y}(n\hat{p}_Y)^k}{k!}$$

$$= 2\bar{s}_b n\hat{p}_Y \exp\left(-n\hat{p}_Y\left(1-\exp\left(-\frac{s_m\hat{p}_X}{\bar{s}_b}\right)\right) - \frac{s_m\hat{p}_X}{\bar{s}_b}\right)$$

where $\hat{p}_X = \hat{p}_m = \hat{p}_f = \sqrt{\frac{\mu_f}{s_f}}$ and $\hat{p}_Y = \min\left(1, \frac{\mu_m}{s_m\hat{p}_X}\right)$

**Fixation of mildly deleterious mutations via hitchhiking**

It is worth comparing the rates at which sheltered alleles fix by hitchhiking relative to the fixation rates of mildly deleterious mutations. Mildly deleterious mutations are expressed in heterozygotes, with typical dominance coefficients of $h \sim 0.25$ (Manna et al. 2011; Charlesworth 2015). For loci mutating to mildly deleterious alleles, we assume that the heterozygous fitness effects are strong relative to the mutation rate ($s_m h, s_f h \gg \mu_m, \mu_f$), in which case the evolutionary dynamics under mutation and selection are well-approximated by:

$$p_m' = p_f(1 - s_m h) + \mu_m$$

$$p_f' = \frac{1}{2}(p_f + p_m)(1 - s_f h) + \mu_f$$

$$p_Y' = p_Y(1 - s_m h) + \mu_m$$

yielding the following equilibria:

$$\hat{p}_f = \frac{\mu_m(1 - s_f h) + 2\mu_f}{2 s_f h + s_m h(1 - s_f h)}$$

$$\hat{p}_m = \hat{p}_f(1 - s_m h) + \mu_m$$

$$\hat{p}_Y = \frac{\mu_m}{s_m h}$$

To distinguish between the mutation rate and fitness effects of mildly deleterious alleles and those of LOF alleles in our sheltering model, let $s_d$ represent the *heterozygous* effect of mildly deleterious mutations in males, and $\mu_d$ represent the mildly deleterious mutation rate per Y-linked locus, so that $\hat{p}_{Y,d} = \mu_m / s_d$ is equilibrium frequency for the locus. Following a similar model by Orr and Kim (1998), we assume that the fitness effects of mildly deleterious alleles are constant across loci, and fitness effects are multiplicative across loci. The fitness associated with a Y chromosome that carries a beneficial mutation and $k$ mildly deleterious alleles is $(1 - s_d)^k(1 + s_b)$, and the probability of establishment for such a haplotype is:

$$\sim 2((1 - s_d)^k(1 + s_b) - 1) \approx 2(s_b - s_d k)$$

which is valid $s_b > s_d k$, and the fixation probability is otherwise zero. Supposing that $s_b$ follows an exponential distribution with mean $\bar{s}_b$, then the fixation probability of a new beneficial mutation that arises in association with $k$ deleterious alleles is:

$$\Pr(\text{fix}|k) \approx \int_{s_d k}^{\infty} 2(s_b - s_d k)\frac{1}{\bar{s}_b} e^{-s_b/\bar{s}_b} ds_b = 2\bar{s}_b \exp\left(-\frac{s_d k}{\bar{s}_b}\right)$$

The number of mildly deleterious mutations segregating on the Y chromosome is Poisson distributed with mean of $L\mu_m/s_d$, where $L$ is the number of Y-linked loci mutating to mildly deleterious alleles. The expected number of mildly deleterious mutations that become fixed for each new beneficial mutation entering the population is:

$$\mathrm{E}(k_d) = \sum_{k=0}^{n} k \Pr(\text{fix}|k) \frac{e^{-L\mu_m/s_d}(L\mu_m/s_d)^k}{k!} \approx 2\bar{s}_b \sum_{k=0}^{n} k \exp\left(-\frac{s_d k}{\bar{s}_b}\right) \frac{e^{-L\mu_m/s_d}(L\mu_m/s_d)^k}{k!}$$

$$= 2\bar{s}_b \frac{U_d}{s_d} \exp\left(-\frac{U_d}{s_d}\left(1 - \exp\left(-\frac{s_d}{\bar{s}_b}\right)\right) - \frac{s_d}{\bar{s}_b}\right)$$

where $U_d = L\mu_m$ is the total Y chromosome mutation rate to mildly deleterious alleles.

For purposes of comparison, let us consider the case where LOF alleles are homozygous lethal ($s_f = s_m = 1$), in which case

$$\mathrm{E}(k_{LOF}) \approx 2\bar{s}_b \frac{U_{LOF}}{\sqrt{\mu_f}} \exp\left(-\frac{U_{LOF}}{\sqrt{\mu_f}}\left(1 - \exp\left(-\frac{\sqrt{\mu_f}}{\bar{s}_b}\right)\right) - \frac{\sqrt{\mu_f}}{\bar{s}_b}\right)$$

where $U_{LOF} = n\mu_m$ is the total Y chromosome mutation rate to LOF alleles. Here, we see that the expressions for mildly deleterious and LOF mutations become identical when $s_d = \sqrt{\mu_f}$. Fixation of homozygous lethal LOF alleles becomes more permissible than mildly deleterious alleles when $s_d > \sqrt{\mu_f}$. Fixation of LOF alleles becomes even more permissible if genes are not essential for males ($s_m < 1$).

**Fixation of LOF alleles via background selection**

We will assume that LOF mutations are recessive, and that a set of background loci are primary selected in heterozygous state. We will assume that background loci are at deterministic mutations-selection balance, and that LOF mutations are initially absent from the Y chromosome. The X-linked LOF allele frequencies are at the mutation-selection-drift equilibrium conditioned on the absence of LOF mutations on the Y.

Following the general approach of Charlesworth (1994) for modelling background selection, the fixation probability of a new LOF mutation arising on the Y will depend on:

- The homozygous fitness cost of the LOF allele in males ($s_m$)
- The census ($N$) effective population size ($N_e$); we assume an equal sex ratio and equal variance in reproductive success
- The initial frequency of new mutations on the Y chromosome ($p = 2/N$)
- The fraction of the Y chromosome that is free of deleterious mutations at background loci ($f_0$, further defined below)

Following the standard theory of mutation-selection balance (Orr 2000), and assuming no LD among the background loci, the frequency of the least loaded Y chromosome class will be:

$$f_0 = e^{-U_Y/s_H}$$

where $U_Y$ is the total Y-linked mutation rate at the background loci and $s_H$ is the harmonic mean heterozygous fitness cost of background mutations.

Because a new LOF will be eliminated if it arises in a loaded background, we focus on the subset that lands on an unloaded background. For those mutations, the initial frequency will be $p_0 = p/f_0$ (Charlesworth 1994). Conditioned on the LOF mutation residing in the least loaded class, the mean and variance in allele frequency change for a LOF mutation at frequency $p_Y$ will be (respectively):

$$M_Y \approx -s_m \bar{p}_X p_Y (1 - p_Y)$$

And

$$V_Y = \frac{2p_Y(1-p_Y)}{f_0 N_e}$$

where $\bar{p}_X$ is the expected X-linked frequency of the LOF allele (Nei 1970; see Appendix 2). Conditioned on the Y-linked LOF mutation landing in the least loaded class, the probability of its fixation will be:

$$u_Y(p_0) = \frac{\exp(f_0 N_e s_m \bar{p}_X p_0) - 1}{\exp(f_0 N_e s_m \bar{p}_X) - 1} = \frac{\exp\left(\frac{2N_e}{N} s_m \bar{p}_X\right) - 1}{\exp(f_0 N_e s_m \bar{p}_X) - 1}$$

Since the probability of a random LOF mutation landing on an unloaded background is $f_0$, the overall fixation probability of a LOF mutation will be:

$$\Pr(\text{LOF fixes}) = f_0 u_Y(p_0) = f_0 \frac{\exp\left(\frac{2N_e}{N} s_m \bar{p}_X\right) - 1}{\exp(f_0 N_e s_m \bar{p}_X) - 1}$$

For point of contrast, we can also calculate the fixation rate of mildly deleterious mutations through background selection. For mutation with heterozygous fitness cost of $s_d$, the probability of fixation under background selection is:

$$\Pr(\text{del. mutation fixes}) = f_0 \frac{\exp\left(\frac{2N_e}{N} s_d\right) - 1}{\exp(f_0 N_e s_d) - 1}$$

The fixation probability for LOF alleles will be higher whenever $s_m \bar{p}_X < s_d$. In the case of lethal mutations in a very large population, this condition simplifies to $\sqrt{\mu_f} < s_d$. The condition becomes more permissive for LOF mutations in non-essential genes.

#### Appendix 4: Fixation probabilities with incomplete recessivity

The stochastic results presented above apply to completely recessive LOF alleles. Here, we extend these results to incomplete recessivity.

***Fixation of X-linked LOF mutations.*** We first consider the fixation of LOF alleles in male-limited genes on the X. The ancestral population is (potentially) segregating for a LOF allele in the Y-linked copy of the gene, and the X-linked copy is initially fixed for the functional allele ($p_f = p_m = 0$ in the ancestral population). With the Y-linked LOF allele segregating at frequency $p_Y$, the fixation probability of a new X-linked LOF allele is:

$$U_X(p_0) = \frac{\int_0^{p_0} G(x)dx}{\int_0^1 G(x)dx} = \frac{\exp\big(N_e s_m\big(h + p_Y(1-2h)\big)p_0\big) - 1}{\exp\Big(N_e s_m\big(h + p_Y(1-2h)\big)\Big) - 1}$$

where:

$$G(x) = \exp\left(-2\int_0^x \frac{M_X}{V_X}dp\right) = \exp\big(N_e s_m\big(h + p_Y(1-2h)\big)x\big)$$

$$M_X = -\frac{1}{3}s_m p(1-p)\big(h + p_Y(1-2h)\big)$$

$$V_X = \frac{2p(1-p)}{3N_e}$$

The initial frequency of a new X-linked mutation is $p_0 = 2/(3N)$, which has a fixation probability of:

$$U_X(p_0 = 2/(3N)) \approx \frac{\frac{2}{3}\frac{N_e}{N}s_m\big(h + p_Y(1-2h)\big)}{\exp\Big(N_e s_m\big(h + p_Y(1-2h)\big)\Big) - 1}$$

In the absence of X-linked LOF alleles, the deterministic dynamics for the Y-linked LOF allele are:

$$\Delta p_Y = \frac{s_m h(1-p_Y)(\hat{p}_Y - p_Y)}{1 - p_Y s_m h}$$

where $\hat{p}_Y = \mu_m/(s_m h)$ is the deterministic mutation-selection balance equilibrium, which is valid when $s_m h > \mu_m$. Representative results are shown in the following figure, in which $\mu_m = 10^{-6}$. Naturally, fixation probabilities for X-linked mutations will decrease even further when there is purifying selection in females in addition to males (i.e., when $s_f > 0$)

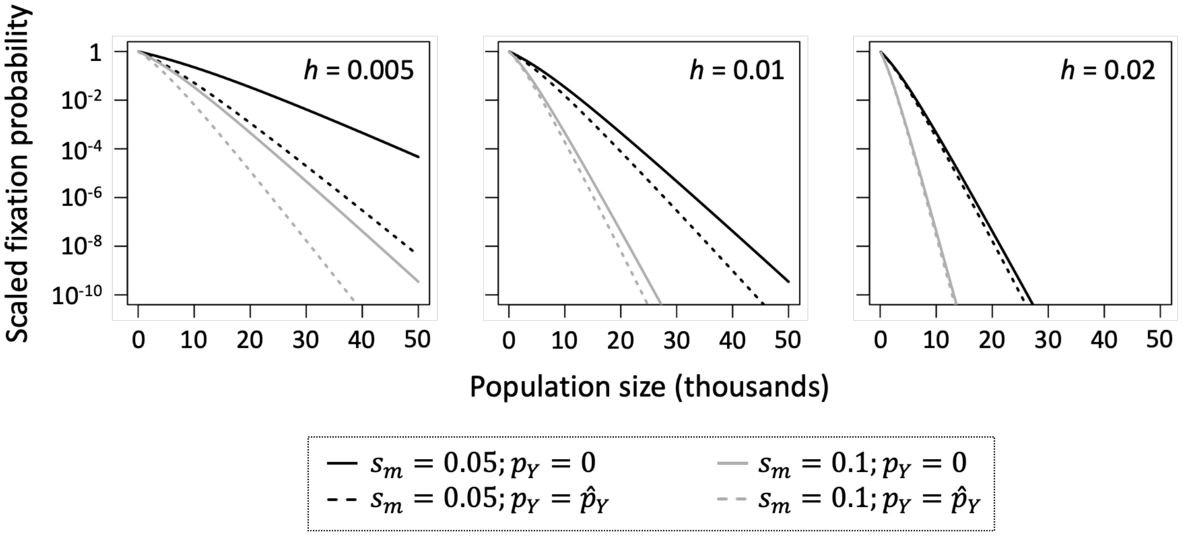


In populations where LOF alleles for gene are not segregating on the Y ($p_Y = 0$), the fixation probability of an X-linked LOF allele will be greater than 1/10 the fixation probability of a neutral variant when $N_e s_m h < \sim 3.6$. Thus, we might consider fixation to be reasonably likely when dominance falls below the critical threshold $h = 3.6/(N_e s_m)$. Thus, while male-essential genes can be deterministically lost from the X when $h = 0$ (see Appendix 1), expression in heterozygotes strongly constrains the loss of such genes unless the population is small.

***Fixation of Y-linked LOF mutations.*** We next consider the fixation probabilities of Y-linked LOF alleles in genes with arbitrary effects in each sex. The deterministic evolutionary dynamics of a Y-linked LOF mutation are described by:

$$\Delta p_Y \approx -s_m p_Y (1 - p_Y)\big(h + p_X(1 - 2h)\big)$$

where $p_X$ is the frequency of the LOF allele on the X chromosome. The fixation probability of a Y-linked mutant becomes:

$$U_Y(p_0) = \frac{\int_0^{p_0} G(x)dx}{\int_0^1 G(x)dx} = \frac{\exp\left(N_e s_m\left(h + p_X(1-2h)\right)p_0\right) - 1}{\exp\left(N_e s_m\left(h + p_X(1-2h)\right)\right) - 1}$$

where:

$$G(x) = \exp\left(-2\int_0^x \frac{M_Y}{V_Y} dp_Y\right) = \exp\left(N_e s_m\left(h + p_X(1-2h)\right)x\right)$$

These expressions assume that the effective population size for the Y chromosome is one-quarter the size of an autosome. The initial frequency of a new Y-linked mutation is $p_0 = 2/N$ has a fixation probability of:

$$U_Y(p_0 = 2/N) \approx \frac{\frac{2N_e}{N} s_m\left(h + p_X(1-2h)\right)}{\exp\left(N_e s_m\left(h + p_X(1-2h)\right)\right) - 1}$$

In absence of segregating LOF alleles on the Y, the deterministic dynamics of the X-linked LOF allele are described by:

$$\Delta p_X \approx -\frac{2}{3} s_f p(1-p)\left(h + p(1-2h)\right) - \frac{1}{3} s_m h p(1-p) + \frac{2\mu_f + \mu_m}{3}(1-p)$$

which has the mutation-selection equilibrium:

$$\hat{p}_X = \frac{-(2s_f + s_m)h + \sqrt{\left((2s_f + s_m)h\right)^2 + 8s_f(1-2h)(2\mu_f + \mu_m)}}{4s_f(1-2h)}$$

Assuming that $N_e s_f$ is sufficiently large that the X-linked mutation remains rare in the population (hence terms of $(1-p_X)^{-1}$ are negligible), then the stationary distribution for the X-linked gene (in the absence of segregating LOF alleles on the Y) is:

$$f(p_X) = C p_X^{N_e(2\mu_f + \mu_m) - 1} \exp\left(-N_e(2s_f + s_m)h p_X - N_e s_f(1-2h)p_X^2\right)$$

where *C* is a constant of integration. The scaled fixation probability, evaluated at the expected allele frequency on the X ($\bar{p}_X = \int p_X f(p_X) dp_X$), is:

$$\frac{U_Y(p_0 = 2/N)}{2/N} \approx \frac{N_e s_m\big(h + \bar{p}_X(1-2h)\big)}{\exp\Big(N_e s_m\big(h + \bar{p}_X(1-2h)\big)\Big) - 1}$$

in which $s_m\big(h + \bar{p}_X(1-2h)\big)$ is the effective strength of purifying selection against Y-linked mutations. The effect of selection in females on the strength of purifying selection against Y-linked alleles can be evaluated by differentiating $s_m\big(h + \bar{p}_X(1-2h)\big)$ with respect to $s_f$, which yields:

$$\frac{ds_m\big(h + \bar{p}_X(1-2h)\big)}{ds_f} \approx s_m \frac{d\bar{p}_X}{ds_f}(1-2h)$$

Since $\frac{d\bar{p}_X}{ds_f}$ is always negative, the strength of selection against partially or completely recessive Y-linked mutations ($h$ < 0.5) is expected to always decrease as the strength in selection in females increases. Consequently, the probability of gene decay from the Y ($U_Y(p_0 = 2/N)$) should increase with increasing $s_f$. Note that the qualitative effect is independent of how important the gene is for males (i.e., the sign of $\frac{ds_m(h+\bar{p}_X(1-2h))}{ds_f}$ does not depend on $s_m$).

If LOF alleles of a gene are not segregating on the X, fixation of a LOF mutation on the Y will be reasonably likely (i.e., the probability of fixation is at least 10% that of a neutral mutation) when $N_e f_0 s_m h < \sim 3.6$, which includes the effect of background selection.

#### Supplementary Figures

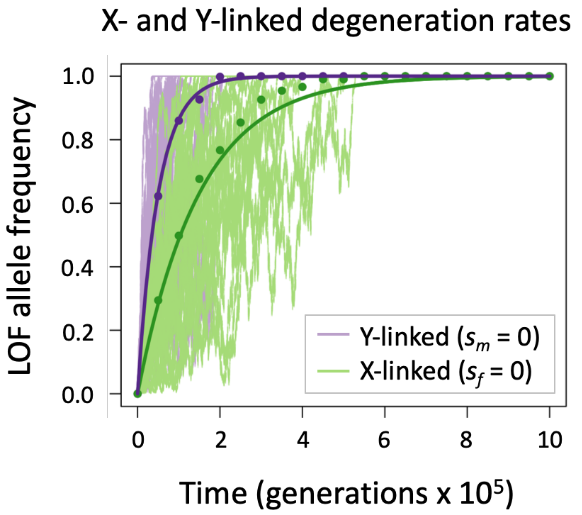


**Figure S1.** Evolutionary dynamics of LOF mutations that eventually become fixed in sex-limited genes: Further results. Degeneration is shown in a smaller population than is presented in Fig. 3b of the main text. Here, the population size is an order of magnitude smaller ($N_e = 10^4$ rather than $N_e = 10^5$), but with all other parameters remaining the same (*i.e.*, $\mu_f = 10^{-5}$, $\mu_m = 2\mu_f$, $s_f = 1$ for Y-linked genes, and $s_m = 1$ for X-linked genes). 30 simulation runs are shown for each chromosome, with individual trajectories (thin, pale lines) scattered about the analytical predictions (bold curves) and circles denoting mean LOF frequencies across the set of simulated trajectories. The average trajectories LOF allele frequency trajectories are highly predictable (i.e., we see good alignment between the bold curves and the circles). However, individual allele frequency trajectories exhibit greater variability in small compared to large populations (se Fig. 3b in the main text for contrast), as indicated by the broad range of frequency states denoted in pale purple (for the Y) and pale green (for the X).

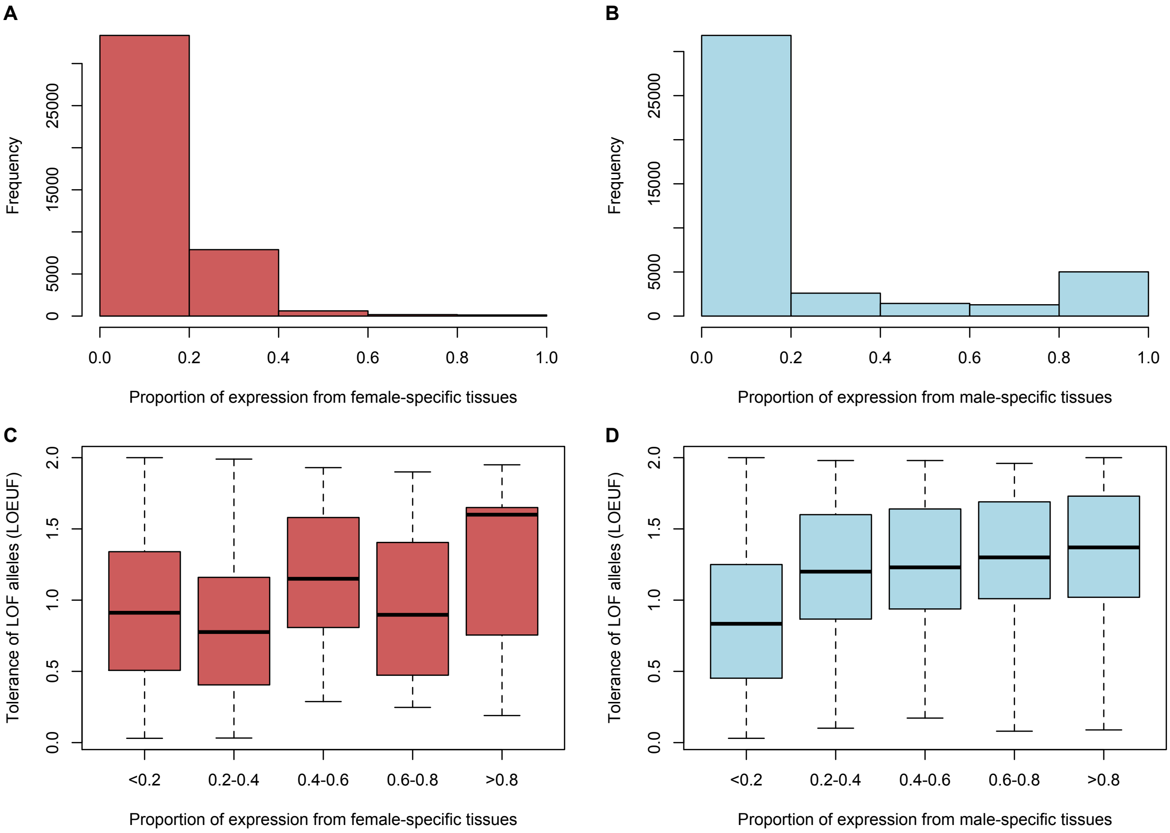


**Figure S2.** Estimates of tolerance of loss of function (LOF) alleles from the gnomAD dataset (version 2.1; see the Supplementary Methods, immediately below). **Panels A and B:** Number of genes per sex-bias category (measured as the proportion of total expression derived from female- and male-specific tissues, respectively). **Panels C and D:** Distribution of LOEUF scores for different categories of female- and male-biased genes. LOEUF scores quantify the abundance of segregating LOF alleles relative to a neutral evolutionary model in which there is no purifying selection against LOF mutations. The LOEUF metric is continuous between zero and two, with LOEUF = 0 representing the strongest degree of constraint (strong purifying selection against LOF alleles) and LOEUF = 2 representing the least constraint.

### Supplementary Methods

#### Survey of sex-biased and sex-specific expression

The following databases were mined for RNA-seq datasets containing whole-body data from both males and females: the NCBI Short Reads Archive (https://www.ncbi.nlm.nih.gov/sra), the NCBI GEO Datasets (https://www.ncbi.nlm.nih.gov/gds), and Google Scholar (https://scholar.google.com). Priority was given to datasets for which both adult and juvenile stages were sampled, or which allowed for broader phylogenetic sampling (brine shrimp, tardigrade, and zebrafish datasets were included despite only containing adult data as their (sub)phylum was otherwise not represented). The list of datasets with information on replicate number, developmental stages that were sampled, and data type, are provided in Supplementary Table S2.

When gene expression values were provided, either as GEO datasets or supplementary tables of the associated manuscripts, these were downloaded and used directly. If expression values were provided as raw read counts, CPM values (counts per million) were estimated before proceeding. FPKM, RKPKM and TPM values were used directly. Only RNA-seq reads were available for zebrafish (Danio rerio) and wood white butterflies (Leptidea sinapis). For these two species, CDS sequences were obtained from Ensembl (Danio_rerio.GRCz11.cds.all.fa) and from the NCBI genome assembly page (GCF_905404315.1_ilLepSina1.1_cds_from_genomic.fna) respectively. Only the longest CDS was kept for each gene, and expression values were obtained using Kallisto (version 0.50.1; Bray et al. 2016).

Gene expression values were quantile-normalized across each dataset with NormalyzerDE (Willforss et al. 2019). For each gene, expression values were averaged by sex. In order to avoid including very low expression genes, which may not be biologically relevant, we removed genes that had summed male and female expression below the 25th percentile. A second round of quantile normalization was performed on the averaged male and female values (as filtering after averaging replicates can lead to uneven distributions despite the initial normalization). In order to avoid removing sex-specific genes, 0.1 was added to all expression values before the log2 ratio of male to female expression was calculated; this ratio was then used to select genes with varying fold-change levels of sex-bias. To infer the number of sex-specific genes, we also calculated the metric male_expression/(male_expression + female_expression), and selected genes for which this metric was below 0.01 (female-specific) or above 0.99 (male-specific).

In the case of humans, no whole-body data was obviously available, and they were therefore not included in the estimation of sex-bias. Instead, median gene expression values for specific tissues (obtained from 4 to many hundreds of individuals depending on the tissue) were downloaded from GTEX:

GTEx_Analysis_2017-06-05_v8_RNASeQCv1.1.9_gene_median_tpm.gct.gz from:
https://www.gtexportal.org/home/downloads/adult-gtex#bulk_tissue_expression

While for most tissues the median expression is derived from male and female individuals, several tissues are sex specific: Uterus, Vagina, Ovary, Ectocervix, Endocervix, Fallopian Tube, and Breast Mammary are female-specific, and Prostate and Testis are male-specific. For each gene, expression was summed across all tissues. Genes with total expression below the 25th percentile were removed, to avoid including very low expression genes. Genes were then classified as male-specific, if over 99% of their expression came from male-specific tissues, and female-specific, if over 99% of their expression came from female-specific tissues.

**Analysis of GNOMAD scores for tolerance of human genes to LOF mutations**

The table "pLoF Metrics by Gene" (GNOMAD V2.1.10; Karczewski et al. 2020) was obtained from https://gnomad.broadinstitute.org/downloads#v2. We focused on this version (which included 141,456 human genomes and exomes) because scores were provided per gene rather than per transcript, making it possible to combine them with the sex-bias estimates obtained in the previous section. As noted on the GNOMAD website, "genes in each LOEUF decile are fairly stable between gnomAD v2.1.1 and v4.0", such that the results shown here should not be version-dependent. The distributions of loss-of-function observed/expected upper bound fraction scores (LOEUF scores, column "oe_lof_upper" in the table) were plotted by category of sex bias, defined based on the proportion of gene expression derived from female- and male-specific tissues (see above).

#### Table S2. Species and datasets included in the expression analysis

| Species | Common name | (Sub)Phylum | Stage | Replicates | individuals per replicate | Notes | Reference* |
|---|---|---|---|---|---|---|---|
| *Paramacrobiotus metropolitanus* | tardigrade | Tardigrada | Not specified but sexually differentiated | 4 Male / 3 Female | 250 | Using TMM values in Geo dataset | [1] |
| *Caenorhabditis brenneri* | roundworm | Nematoda | L4+adult | 3 per sex | >400 | Using RSEM expression values in Geo dataset (FPKM) | [2] |
| *Caenorhabditis remanei* | roundworm | Nematoda | L4+adult | 3 per sex | >400 | Using RSEM expression values in Geo dataset (FPKM) | [2] |
| *Caenorhabditis japonica* | roundworm | Nematoda | L4+adult | 3 per sex | >400 | Using RSEM expression values in Geo dataset (FPKM) | [2] |
| *Danio rerio* | zebrafish | Vertebrata | Adult | 2 per sex | 1 | Restimated TPM from raw RNA-seq reads | [3] |
| *Artemia sinica* | brine shrimp | Arthropoda | Adult | 2 per sex | 1 | Using TPM values provided in the supplementary materials. | [4] |
| *Homo sapiens* | human | Vertebrata | Adults | at least 4 per tissue | 1 | Using median expression per gene from version 6 of GTEX | [5] |
| *Drosophila melanogaster* | fruit fly | Arthropoda | L3 + pupae + adults | 3 per stage / sex | 1 | Counts per million mapped reads calculated using read counts provided in supplementary material; original data from https://doi.org/10.1101/034728 | [6] |
| *Timema californicum* | Stick insect | Arthropoda | Hatchlings + juveniles + adults | at least 3 per stage / sex | 1 | Counts per million mapped reads calculated using read counts provided in supplementary material | [6] |
| *Schistosoma japonicum* | blood fluke | Platyhelminthes | 14 to 28 days (sexual maturation) | 3 per time point / sex | >80 | Using TPM table provided in Elkrewi et al 2021; original data from various datasets | [7] |
| *Schistosoma mansoni* | blood fluke | Platyhelminthes | cercariae + somulae + adults | at least 2 per stage / sex | 1 to >1000 | Using TPM table provided in Elkrewi et al 2021; original data from various datasets | [7] |
| *Leptidea sinapis* | wood white (butterfly) | Arthropoda | L5+pupae+adults | 4 to 6 per stage / sex | 1 | Restimated TPM from raw RNA-seq reads(NCBI bioproject PRJEB24745); only long-day samples used as they contained all developmental stages. | [8] |

*** Expression data references:**

[1] Sugiura K, Yoshida Y, Hayashi K, Arakawa K, Kunieda T, Matsumoto M. 2024. Sexual dimorphism in the tardigrade *Paramacrobiotus metropolitanus*. *Zoological Letters* 10:11.
[2] Thomas CG, Li R, Smith HE, Woodruff GC, Oliver B, Haag ES. 2012. Simplification and desexualization of gene expression in self-fertile nematodes. *Current Biology* 22:2167-2172.
[3] King AC, Zenker AK. 2023. Sex blind: bridging the gap between drug exposure and sex-related gene expression in Danio rerio using next-generation sequencing (NGS) data and a literature review to find the missing links in pharmaceutical and environmental toxicology. *Frontiers in Toxicology* 5:1187302
[4] Elkrewi M, Khauratovich U, Toups MA, Bett VK, Mrnjavac A, Macon A, Fraisse C, Sax L, Huylmans AK, Hontoria F, Vicoso B. 2023. ZW sex-chromosome evolution and contagious parthenogenesis in Artemia brine shrimp. *Genetics* 222:iyac123.
[5] Lonsdale J, et al. 2013. The genotype-tissue expression (GTEx) project. *Nature Genetics* 45:580-585.
[6] Djordjevic J, Dumas Z, Robinson-Rechavi M, Schwander T, Parker DJ. 2022. Dynamics of sex-biased gene expression during development in the stick insect Timema californicum. *Heredity* 129:113-122.
[7] Elkrewi M, Moldovan MA, Picard MAL, Vicoso B. 2021. Schistosome W-linked genes inform temporal dynamics of sex chromosome evolution and suggest candidate for sex determination. *Mol Biol Evol*. 38:5345-5358.
[8] Leal L, Talla V, Källman T, Friberg M, Wiklund C, Dincă V, Vila R, Backström N. 2018. Gene expression profiling across ontogenetic stages in the wood white (*Leptidea sinapis*) reveals pathways linked to butterfly diapause regulation. *Molecular Ecology* 27:935-948.
